# Targeted *in situ* cross-linking mass spectrometry and integrative modeling reveal the architectures of Nsp1, Nsp2, and Nucleocapsid proteins from SARS-CoV-2

**DOI:** 10.1101/2021.02.04.429751

**Authors:** Moriya Slavin, Joanna Zamel, Keren Zohar, Siona Eliyahu, Merav Braitbard, Esther Brielle, Leah Baraz, Miri Stolovich-Rain, Ahuva Friedman, Dana G Wolf, Alexander Rouvinski, Michal Linial, Dina Schneidman-Duhovny, Nir Kalisman

**Affiliations:** Department of Biological Chemistry, Institute of Life Sciences, The Hebrew University of Jerusalem, Jerusalem, Israel; Hadassah Academic College Jerusalem, Jerusalem, Israel; Department of Microbiology and Molecular Genetics, Institute for Medical Research Israel-Canada (IMRIC), The Kuvin Center for the Study of Infectious and Tropical Diseases, The Hebrew University-Hadassah Medical School, The Hebrew University of Jerusalem, Jerusalem, Israel; Clinical Virology Unit, Hadassah Hebrew University Medical Center, Jerusalem, Israel; The Rachel and Selim Benin School of Computer Science and Engineering, The Hebrew University of Jerusalem, Jerusalem, Israel

## Abstract

Atomic structures of several proteins from the coronavirus family are still partial or unavailable. A possible reason for this gap is the instability of these proteins outside of the cellular context, thereby prompting the use of in-cell approaches. *In situ* cross-linking and mass spectrometry (*in situ* CLMS) can provide information on the structures of such proteins as they occur in the intact cell. Here, we applied targeted *in situ* CLMS to structurally probe Nsp1, Nsp2, and Nucleocapsid (N) proteins from SARS-CoV-2, and obtained cross-link sets with an average density of one cross-link per twenty residues. We then employed integrative modeling that computationally combined the cross-linking data with domain structures to determine full-length atomic models. For the Nsp2, the cross-links report on a complex topology with long-range interactions. Integrative modeling with structural prediction of individual domains by the AlphaFold2 system allowed us to generate a single consistent all-atom model of the full-length Nsp2. The model reveals three putative metal binding sites, and suggests a role for Nsp2 in zinc regulation within the replication-transcription complex. For the N protein, we identified multiple intra- and inter-domain cross-links. Our integrative model of the N dimer demonstrates that it can accommodate three single RNA strands simultaneously, both stereochemically and electrostatically. For the Nsp1, cross-links with the 40S ribosome were highly consistent with recent cryo-EM structures. These results highlight the importance of cellular context for the structural probing of recalcitrant proteins and demonstrate the effectiveness of targeted *in situ* CLMS and integrative modeling.

## Introduction

The genome of SARS-CoV-2 encodes 29 major proteins - Sixteen non-structural proteins (Nsp1-16), four structural proteins (S, E, M, N), nine major Orfs, and several additional non-canonical gene products^1^. Once the human cell is infected, the viral proteins are engaged in a network of protein-protein interactions (PPI) that lead to alterations in signalling pathways and to a global shift towards viral protein production^2,3^. Despite significant progress in viral protein structure determination^4^, there are still gaps in the structural knowledge of several proteins from the coronavirus family^5^. For example, no structure is yet available for Nsp2, despite bioinformatics evidence that predicts most of its sequence to be structured. Another example is the Nucleocapsid protein, which is known to form multi-subunit assemblies that were not yet resolved structurally. A possible reason for these difficulties may be the instability of certain viral proteins in the *in vitro* state. In such cases, purification procedures that are an integral part of mainstream structural approaches (x-ray crystallography, NMR, and cryo-EM) may cause the purified proteins to disassemble, denature, or aggregate. To avoid such artefacts, *in situ* techniques for structural studies are required.

*In situ* cross-linking and mass spectrometry (*in situ* CLMS) allows probing protein structure inside intact cells^6,7^. In this approach, cells are incubated with a membrane-permeable cross-linking reagent, which reacts with the cellular proteins in their native environment. Following the chemical cross-linking, the cells are lysed and their protein content is analyzed by mass spectrometry (MS). Computational search can then identify from the mass spectrometry data the pairs of residues that were covalently linked. Because a link between two residues reports on their structural proximity, the list of identified links is a rich resource for modeling protein structures and interactions^8–12^. *In situ* CLMS had progressed significantly in recent years, with successful applications on isolated organelles^13–17^, bacteria^18,19^, human cells^20–22^, and heart tissue^23^.

An inherent difficulty of *in situ* CLMS is the high complexity of the initial samples that contain the entire cell proteome. Two general strategies have been employed to reduce the complexity prior to the mass spectrometry analysis. One strategy enriches the cross-linked peptides out of the total tryptic digest by either tagging the cross-linker itself^23^ or by extensive chromatography^18,19^.

The other strategy is based on the ability to purify a specific protein-of-interest out of the cell lysate prior to digestion. Wang *et al*.^20,21^ effectively used the second strategy to study the human proteasome by expressing several of its subunits with a biotin tag. We propose the term “targeted *in situ* CLMS” to describe the latter approach, which allows the user to focus the mass spectrometry resources on a small set of predetermined proteins (targets).

In this work we used targeted *in situ* CLMS and integrative modeling^24,25^ to probe the structures of three SARS-CoV-2 proteins: Nsp1, Nsp2, and the Nucleocapsid protein (N). Our motivation for choosing these proteins is the incomplete available knowledge on their structures and functions. Following cell transfection with the tagged protein, we were able to identify considerable cross-link sets of *in situ* origin. Computational integration of the cross-links with additional structural information allowed us to build almost complete models for Nsp2 and the N protein.

## Results

### Targeted *in situ* CLMS to study viral proteins

We employed a targeted strategy for *in situ* CLMS of viral proteins inside intact human cells. To that end, HEK293 cells were transfected with a plasmid of a selected viral protein fused to a Strep tag (**Figure 1**). We then cross-linked the intact cells with a membrane-permeable cross-linker (either DSS or formaldehyde), washed away the excess cross-linker, lysed the cells, and purified the viral protein via the Strep tag. The purification step greatly enriches and simplifies the sample for mass spectrometry, and increases the subsequent identification rate of cross-links on that proteIn.

**Figure 1.**
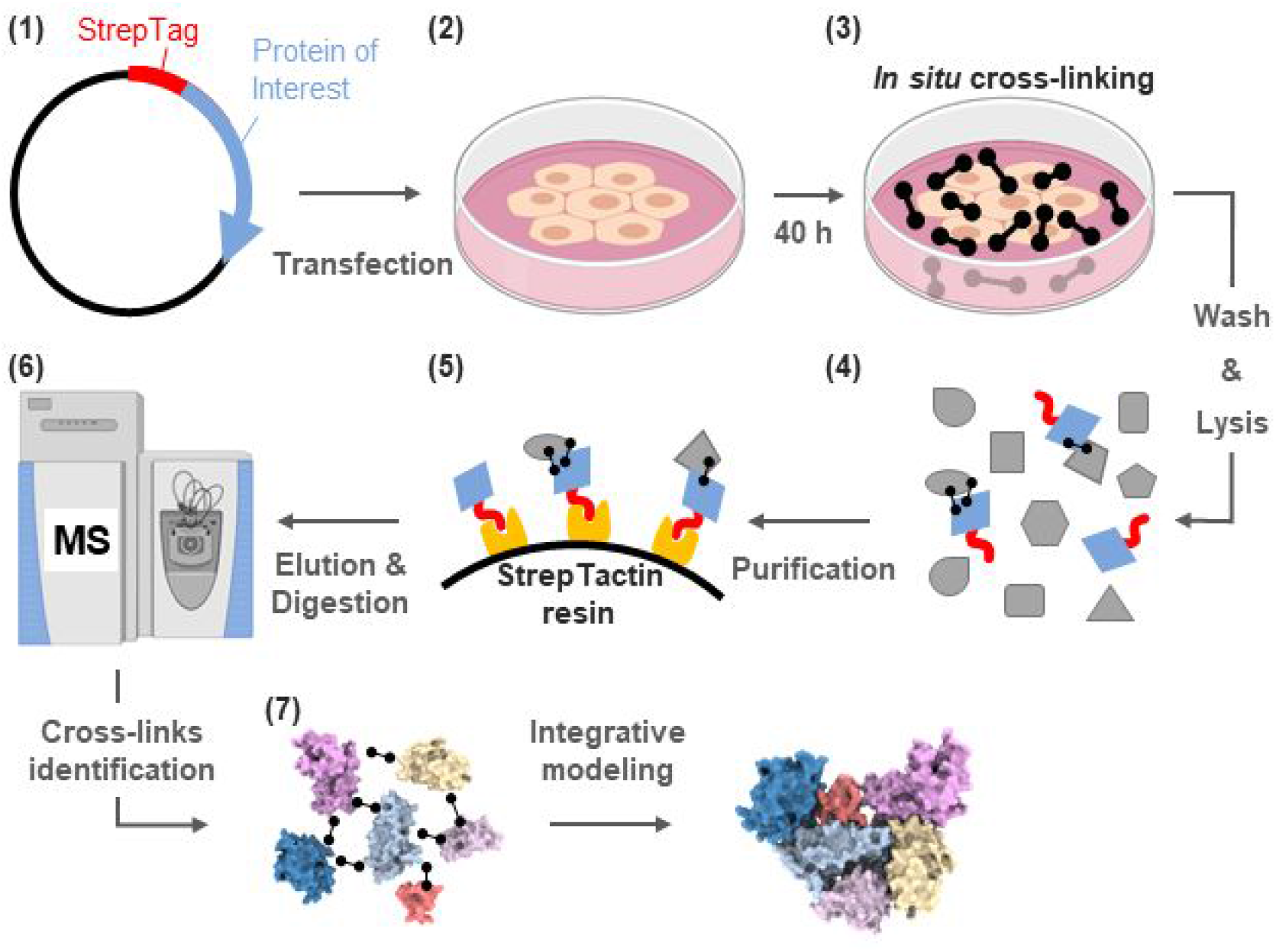
The targeted *in situ* CLMS workflow for a specific protein of interest. **(1)** Cloning of a plasmid for constitutive expression of the viral protein with a Strep tag fused at one of the termini. **(2)** Transfection of human cells in culture with the plasmid, followed by 40 hours of expression. **(3)** The cells are cross-linked *in situ* by a membrane permeable reagent (DSS or formaldehyde, black barbells). **(4)** Washing out the cross-linker before lysis ensures that all the cross-links are of *in situ* origin. **(5)** The protein of interest is purified from the lysate by StrepTactin resin. Cross-linked interactors may co-purify. **(6)** Mass spectrometry (MS) analyses of the purified proteins reveal the protein composition and identify cross-links. **(7)** Integrative modeling generates assemblies using domain structural models and cross-links.

We focused on three SARS-CoV-2 proteins for which the structural information is either missing or incomplete: Nsp1 (180aa), Nsp2 (639aa), and the Nucleocapsid protein (N protein, 419aa). The expression levels of all three proteins peaked 40 hours after transfection. Standard proteomics analyses at peak expression (repeated experiments) detected that: The N protein was the most abundant in the cell, Nsp2 was detected among the 15-30 most abundant proteins, and Nsp1 was among the 250-400 most abundant proteins. The cell morphologies appeared normal, but the adherence of the Nsp1-expressing cells to the plate was considerably weaker. These observations are in accordance with the known toxic role of Nsp1, which is mediated through a global translation inhibition^26^. Following the purifications, proteomics analyses of the elutions detected the tagged proteins to be the most abundant by a large margin for all three proteins and for both the DSS and formaldehyde cross-linking reagents. We conclude that the Strep tag activity is largely unaffected by amine-reactive cross-linking, and mark it as a tag of choice for these pursuits.

While this methodology is similar to the one introduced by Wang *et al*.^20,21^, the current protocol was modified in three major aspects: (i) The incubation time of the cells with the cross-linker was shortened to 20 minutes, rather than 60 minutes in the original protocol. (ii) We used a Strep tag rather than a biotin tag, which allowed us to remove biotinylated proteins from the purification. (iii) We established a transient transfection protocol rather than producing a stable cell line. The transient transfection provides the flexibility of expressing proteins that might be toxic to the cells, such in the case of Nsp1 expression.

### Integrative modeling of Nsp2 based on in situ cross-links

The role of Nsp2 in the viral pathogenicity is poorly understood. Nsp2 is dispensable for viral replication of SARS-CoV in cell culture, although its deletion attenuates viral growth^27^. In infected cells, Nsp2 translocates to the double-membrane vesicles (DMVs) in which the replication-transcription complexes (RTCs) are anchored^28^. It is yet unclear what is the function of Nsp2 in the context of the DMVs. Secondary structure prediction tools^29^ predict that nearly all the sequence of Nsp2 is structured. Yet, this structure remains unknown, thereby making Nsp2 an attractive objective for targeted *in situ* CLMS.

We targeted Nsp2 for *in situ* CLMS and analyzed the resulting mass spectrometry data for proteomics and cross-links. Proteomics analysis revealed ten proteins that co-purify with Nsp2 in significant amounts (**Figure S1A**). Most of these proteins are part of the Prohibitin complex, which was previously shown to interact with the Nsp2 of SARS-CoV^30^. For identification of DSS cross-links, we ran an exhaustive-mode search of the mass spectrometry data against a sequence database comprising the co-purifying proteins and Nsp2. We compiled two non-overlapping cross-link sets from the search results. The primary set (high quality) comprised 43 internal cross-links within Nsp2 and 10 cross-links within the Prohibitin complex (**Figure 2A, Table S1**). The secondary set (medium quality) comprised 38 cross-links, which were mostly within Nsp2 (**Figure 2B, Table S2**). The false detection rate (FDR) for both sets was estimated to be 3% according to a decoy analysis (Methods, **Figures S2**,**S3**). The secondary set only contained cross-links in which one of the peptides was short (4-6 residues). Such short peptides perform poorly in MS/MS fragmentation, and a common practice in most CLMS studies is to ignore them. However, we found that once a suitable filtration of the search results is applied, many of these cross-links have low FDR values. The primary set is used for structure modeling, while the secondary set is used for structure validation. In addition to the two sets of DSS cross-links, we also identified 5 cross-links within Nsp2 from formaldehyde cross-linking **(Figure 2A, Table S1 bottom**) at a FDR of less than 5% according to a decoy analysis with reverse sequences^22^.

**Figure 2.**
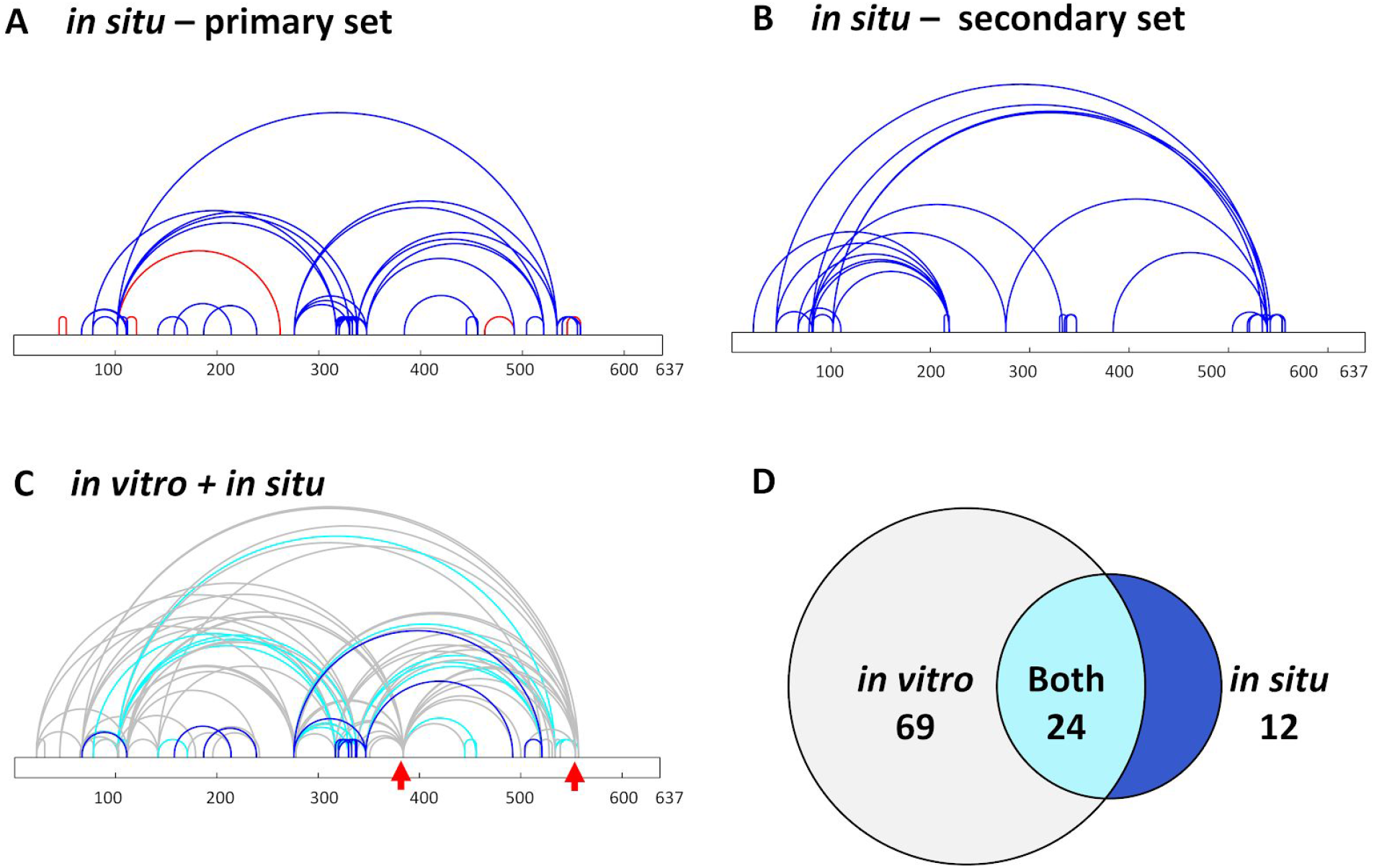
Nsp2 cross-links. **(A)** In situ cross-links depicted as arcs on the sequence of Nsp2. Blue and red arcs represent DSS and formaldehyde cross-links, respectively. **(B)** A secondary set of in situ cross-links that were not part of the primary set. The set comprises cross-links in which one of the peptides is short and has poor fragmentation. The secondary set is only used for final selection of models that are otherwise built by restraints from the primary set. The false detection rate for both sets is 3%. **(C)** Comparison of cross-links identified within Nsp2 from in situ CLMS and in vitro CLMS experiments. Cross-link color corresponds to whether it was identified only in the in situ set (blue), only in the in vitro set (grey), or in both sets (cyan). Note several lysine residues (red arrows) that promiscuously link to multiple sites along the sequence only in the in vitro set. **(D)** The overlap at the residue-pair level between in situ and in vitro cross-link sets.

We attempted to augment the cross-link set by *in vitro* cross-linking of purified Nsp2. To that end, we first purified Nsp2 in HEPES buffer (50mM HEPES pH=8.0, 150mM NaCl), and then cross-linked the purified protein with 1 mM of the soluble BS3 reagent. Because BS3 and DSS share the same cross-linking chemistry, one can directly compare the *in situ* and *in vitro* cross-linked sets (**Figures 2C**,**2D, Table S3**). Surprisingly, the two sets are considerably different, and a significant number of the *in situ* cross-links do not form *in vitro* and vice versa. Notably, several lysine residues link to multiple sites over the entire sequence only in the *in vitro* set. *In vitro* cross-linking with the DMTMM reagent also showed the same promiscuous linking pattern (**Figure S4**) that is not seen in the *in situ* set. These differences are best explained by the tendency of the purified Nsp2 protein to denature and aggregate outside of the cellular context. The inconsistancy between the *in situ* and *in vitro* experiments highlights the crucial information that can be extracted by CLMS from intact cells. It may also underlie the reason for the lack of an atomic full-length structure of Nsp2 to date.

There are no close homologs with solved structure available for Nsp2. We have therefore referred to the Nsp2 model generated by AlphaFold2 from DeepMind^31^. AlphaFold2 has been highly successful in the recent CASP14 round, submitting highly accurate models. The initial AlphaFold2 model violated 17 out of 44 (39%) and 9 out of 29 (31%) cross-links in the primary and secondary sets, respectively. A cross-link is considered violated if the Cα-Cα distance is higher than 25 Å. The violated cross-links were mostly inter-domain ones, while almost all the intra-domain cross-links were satisfied (**Figure 3A**,**B**). To obtain a model consistent with the cross-link set, we divided the AlphaFold2 model into domains (residues 1-104, 105-132, 133-275, 276-345, 512-638). One domain that was not covered by the initial AlphaFold2 model (residues 359-511), was modeled by homology to partial Nsp2 structure of the infectious bronchitis virus (PDB 3ld1^32^, sequence identity 13%). With the availability of the structures for the individual domains, the modeling task is converted into a domain assembly problem. To this end, we applied the CombDock algorithm for multi-molecular assembly based on pairwise docking^33–35^. The six domains served as an input (**Figure 3C**) along with the primary set of cross-links and domain connectivity constraints. We have obtained 62 models that satisfied all the primary set inter-domain cross-links (**Figure 3D**,**E**) with precision of 8 Å. We validated these models with the secondary cross-link set and found nine models that satisfied all but three secondary set cross-links that were in the 25-30 Å range. These models converged into a single cluster with a higher precision of 1 Å (**Figure 3F**).

**Figure 3.**
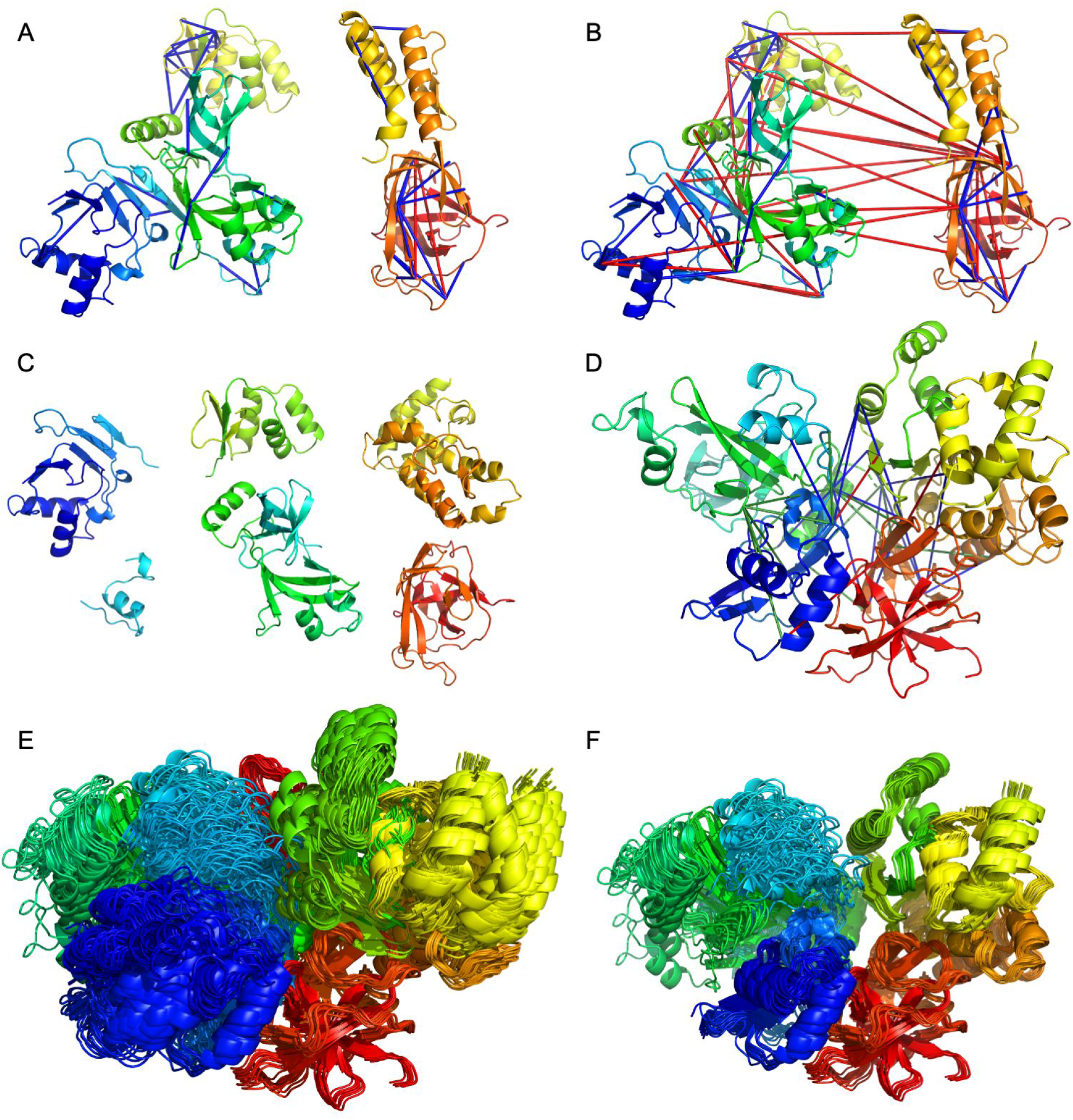
Integrative modeling of Nsp2. The modeling is based on cross-links (satisfied cross-links are in blue, unsatisfied in red). **(A)** Intra-domain cross-links mapped onto the AlphaFold2 model. **(B)** All cross-links mapped onto the AlphaFold2 model. **(C)** Six domains that were given as an input to the modeling pipeline. **(D)** Representative result of the integrative modeling with inter-domain cross-links (blue - primary set, green - secondary set, red - violated cross-links). **(E)** All models that satisfied the primary set inter-domain cross-links. **(F)** Best scoring models according to the secondary cross-link set.

The left view is the same as in Panel B. The large acidic patch in the right view is conserved across human coronaviruses (**Figure S6**).

To infer possible functions of Nsp2, we mapped residues that are fully conserved in all human coronaviruses onto the integrative model. A cluster of conserved cysteine residues around Cys146 prompted a search for possible metal binding sites by studying the distribution of cysteine and histidine residues in the model. We identified three putative metal binding site regions (**Figure 4**). Site 2 is conserved in all coronaviruses, while the other sites occur only in the SARS subfamily. All three sites are solvent accessible within the context of the full integrative model. A search for structures in the Protein Data Bank (PDB) with structurally similar domains (DALI server^36^, Z>3.0) identified zinc binding proteins. Therefore, zinc is the likely ion substrate of Nsp2 as well. An additional histidine residue is located next to the quad of metal-binding residues in all three sites. A possible role for this histidine residue is to modulate the metal binding in a pH-dependent manner. We also note that around the third site there is a clustering of additional cysteine and histidine residues that comprise an incomplete adjacent site. This clustering occurs in SARS-CoV-2, but not in SARS-CoV.

**Figure 4.**
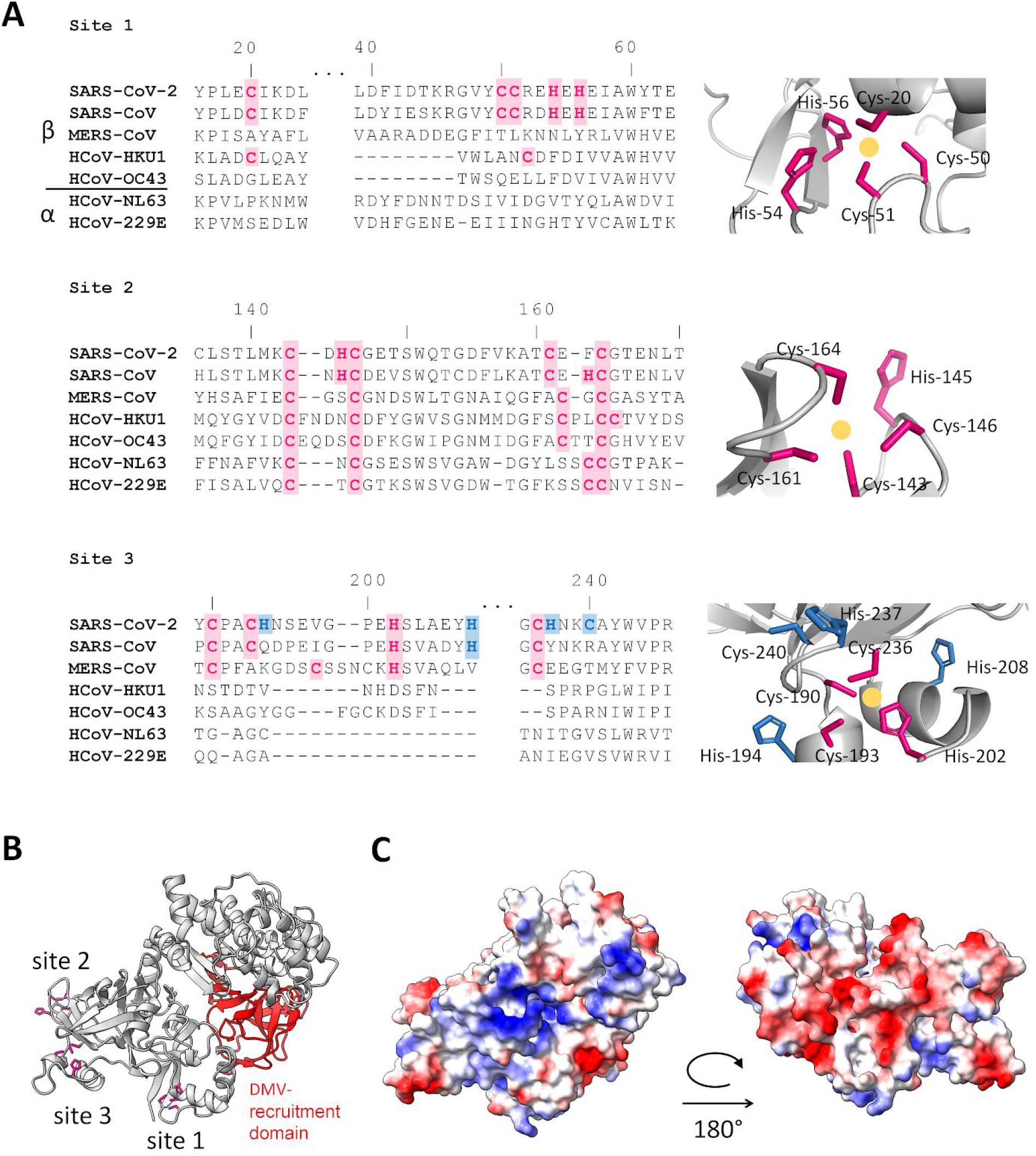
Functional annotation of the Nsp2 model. (**A)** Three putative metal binding sites were identified in the model. Site 2 is conserved in all coronaviruses, while sites 1 and 3 are only occurring in the SARS subfamily. Sidechains of highlighted residues are shown in stick representation on the right. A gold circle marks the putative location of the metal ion. Zinc is the most likely substrate based on structural similarity. In SARS-CoV-2 only, an incomplete site is apparent next to site 3 (blue highlight). (**B)** The three sites are located at the opposite end of the model relative to the domain that recruits the Nsp2 to the replication-transcription complex^28^. (**C)** Electrostatics analysis of the surface of the full model reveals two sides of opposite charges.

The overall evolutionary picture is that of accelerated accumulation of metal binding sites within the SARS subfamily. In coronaviruses, Nsp2 is the least conserved member of the non-structural proteins. Presumably, the reduced evolutionary pressure makes it more amenable to such rapid gain of function. Based on these observations we suggest that Nsp2 plays a role in regulation of zinc levels at the RTCs. Zinc is essential for RNA replication and may be depleted at the RTCs, especially when they are enveloped in the DMVs. The recruitment of Nsp2 to the RTCs therefore establishes a reservoir of zinc that can be exchanged with its surrounding. Finally, we note that zinc regulation is almost certainly not the only function of Nsp2. Analysis of sequence conservation on the surface of the integrative model (**Table S5, Figure S5**) reveals other prominent features that we cannot at present annotate functionally. These include clusters of conserved residues around Pro245 and Tyr619, and a large conserved acidic patch (**Figure 4C, Figure S6**). Future *in situ* CLMS studies in the context of full viral infection will shed more light on these yet uncharacterized functions.

### Assembly of N protein domains based on *in situ* cross-links

The role of the nucleocapsid (N) protein is to pack the viral RNA inside the mature virion. N consists of two domains (RNA binding and dimerization) with known atomic structures for each domain separately^37,38^. These domains are flanked by long linker regions, which are predicted to be largely unstructured (**Figure 5A**), complicating structural determination of the full-length dimer. Recent studies revealed that upon RNA binding N forms higher-order oligomers for tighter packing of the viral RNA inside the capsid^39–42^. Yet, the structural details of this oligomerization are still unknown.

**Figure 5.**
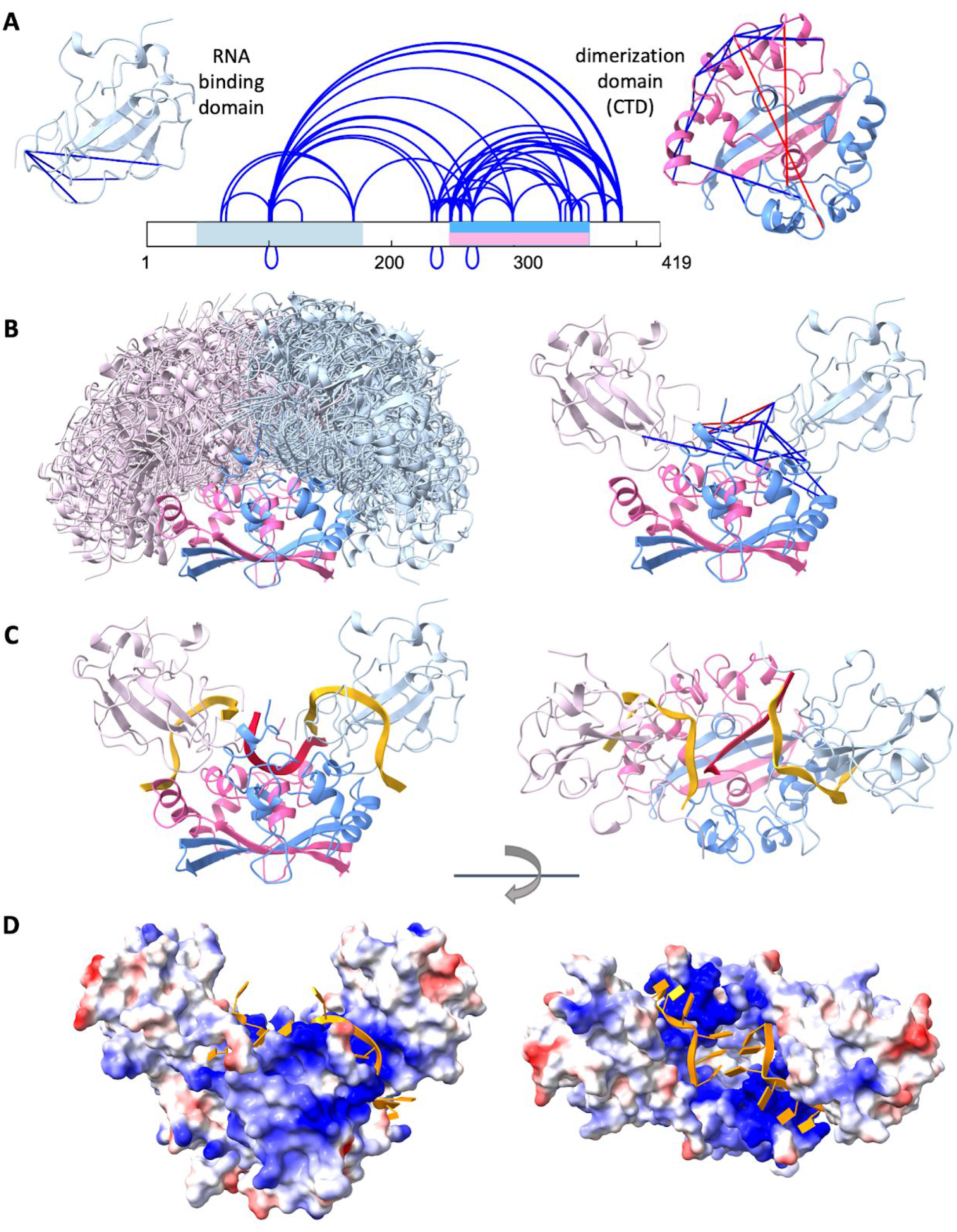
Integrative modeling of the N protein. **(A)** *In situ* cross-links identified within the N protein. The two structured domains of the protein are marked on the sequence in light blue (RNA binding domain) and pink/blue (dimerization domain). The intra-domain cross-links (satisfied cross-links are in blue, unsatisfied in red) are mapped onto the corresponding structures (PDBs 7act^37^ and 6zco^38^ for RNA binding and dimerization domain, respectively). **(B)** Best scoring docking models that satisfy inter-domain cross-links (left) and a representative model with cross-links (right). **(C)** RNA fragments bound to dimerization domain (red) and RNA binding domains (gold). **(D)** The surface of the N protein colored according to the electrostatics potential with two RNA fragments bound to each of the two RNA binding domains in gold color.

Identification of *in situ* DSS cross-links of N (**Figure 5A, Table S6**) followed the same procedure as for Nsp2. We identified 85 unique peptide pairs corresponding to 61 unique cross-linked residue pairs within N, and only one cross-link between N and HNRPU, with FDR of 3% (**Figures S2C and S3C**). This inter-protein cross-link has a borderline fragmentation score and may in fact be a false positive. The cross-links within N can be divided into three groups. The first group contains 19 intra-domain cross-links (4 map to the RNA binding domain and 15 to the dimerization domain). Most of these cross-links fit well with the experimental atomic structures (Cα-Cα distance < 25 Å), three cross-links are in the 26-30 Å range, and two are violated: 266-266 (42 Å) and 248-249 (30.6 Å) (**Figure 5A**). These two cross-links most likely belong to higher order assemblies of the N dimers^39–41,43^. The second group contains the 14 inter-domain cross-links, indicating that the RNA binding and dimerization domains directly interact with each other. The third group contains 27 cross-links between the dimerization domain and the linkers, including a few inter-linker cross-links.

In order to obtain a model of the full N dimer, we performed computational docking^34^ of the dimerization domain (in the dimer form) and two RNA binding domains. All three docked components contained short single RNA strands to ensure that the final model is consistent with the paths of bound RNA. The RNA binding domain and its bound RNA 10-mer were taken from a recent NMR study (PDB 7act^37^). The RNA 6-mer on the dimerization domain (PDB 6zco^38^) was initially docked into the basic groove between the monomers and then refined by a Molecular Dynamics simulation (Methods). The docking was guided by distance restraints derived from the 14 identified inter-domain cross-links^35^. We obtained a single large cluster of models satisfying all the cross-links within 25 Å, except one (residues 100 to 102) which was within 28 Å (**Figure 5B**,**C**). This cross-link is between two lysines located on a flexible loop, therefore the distance can vary. The RNA binding domain binds to the dimerization domain at a well-defined region that was largely shared by all the models. This region comprises residues 247-261 from one chain and residues 296-307 and 343-352 from the other chain.

The integrative model demonstrates that the N dimer can accommodate three RNA single strands simultaneously. Stereochemically, the two RNA binding domains are located far enough from each other to allow a middle RNA single strand to stretch on the basic surface of the dimerization domain without hindrance. We note however that a rearrangement of residues 247-252 in the dimerization domain is required (compared to current crystal structures) in order to allow the entry and exit of that middle strand. Electrostatically, the closest approach of the phosphate backbones between the middle and the side strands is ∼10 Å, which is comparable to proximities observed in the eukaryotic nucleosome^44^. Moreover, the cross-links data suggest that the electrostatic repulsion at these closest-approach regions may be further mitigated by positively charged inter-domain linker regions (see ahead). Overall, the model points to an efficient utilization of the RNA binding capacity of the N dimer, which is required for the packing of the relatively large viral genome.

Of special interest are cross-links that contain an overlapping peptide pair. Such cross-links necessarily report on a direct interaction between two chains of N. We identified four sequence regions that form such cross-links (residues 100-102, 237-237, 248-249, and 266-266). These identifications are supported by well-annotated MS/MS fragmentation spectra (**Figure S8**). In several cases, these identifications are also supported by more than one peptide pair. The dimer model can explain the inter-chain cross-links between Lys100 and Lys102 that are closer than 25 Å in several docking solutions. The inter-chain cross-links of 248-249 and 266-266 are not consistent even with the available crystal structures of the dimerization domain. Most likely they represent a higher-order oligomeric state of N, which we did not try to model here due to insufficient data. Finally, the inter-chain interaction around residue 237 is supported by several cross-links: three different peptide pairs reporting a cross-link between Lys233 and Lys237, and two different peptide pairs reporting a cross-link between Lys237 and Lys237. The multiple cross-links report on a strong interaction of the linker regions that immediately precede the N-terminal of dimerization domain. In the dimer context, these interactions occur on top of the middle RNA strand and between the two other strands bound to the RNA binding domains. This implies that additional positive charges from these linker regions are located between the strands, thereby mitigating their electrostatic repulsion. The same interaction can also serve to clamp the RNA in place.

### Targeted *in situ* CLMS of Nsp1

Nsp1 inhibits host protein translation, thereby interfering with the cellular antiviral response^26^. This short protein comprises a structured N-terminal domain (residues 1-125) and a disordered C-terminal tail (residues 126-180). Two recent cryo-EM studies^46,47^ revealed the C-terminal (residues 148-180) to bind strongly to the mRNA entry tunnel of the ribosome, thus obstructing the tunnel and inhibiting protein synthesis. The results we obtained from targeted *in situ* CLMS of Nsp1 fully support these findings. The main proteins that co-purified with Nsp1 were components of the 40S ribosomal subunit, in particular the ribosomal S3 protein and the eukaryotic translation initiation factor 3 (eIF3) (**Figure S1C**). A search for cross-links among these sequences identified 12 intra-protein cross-links in Nsp1 and two cross-links between Nsp1 and RS3 (**Figure 6A, Table S7**). Another three cross-links were identified within the eIF3 consistent with Nsp1 binding also to the 43S pre-initiation complex^47^. The estimated FDR of this set is less than 5% (**Figures S2D**,**S3D**). The intra-protein cross-links within Nsp1 fit well to the available crystal structure of the N-terminal domain (**Figure 6B**). The two cross-links between Nsp1 and RS3 are in accord with the likely path of the C-terminal along the mRNA entry tunnel (**Figure 6C**).

**Figure 6.**
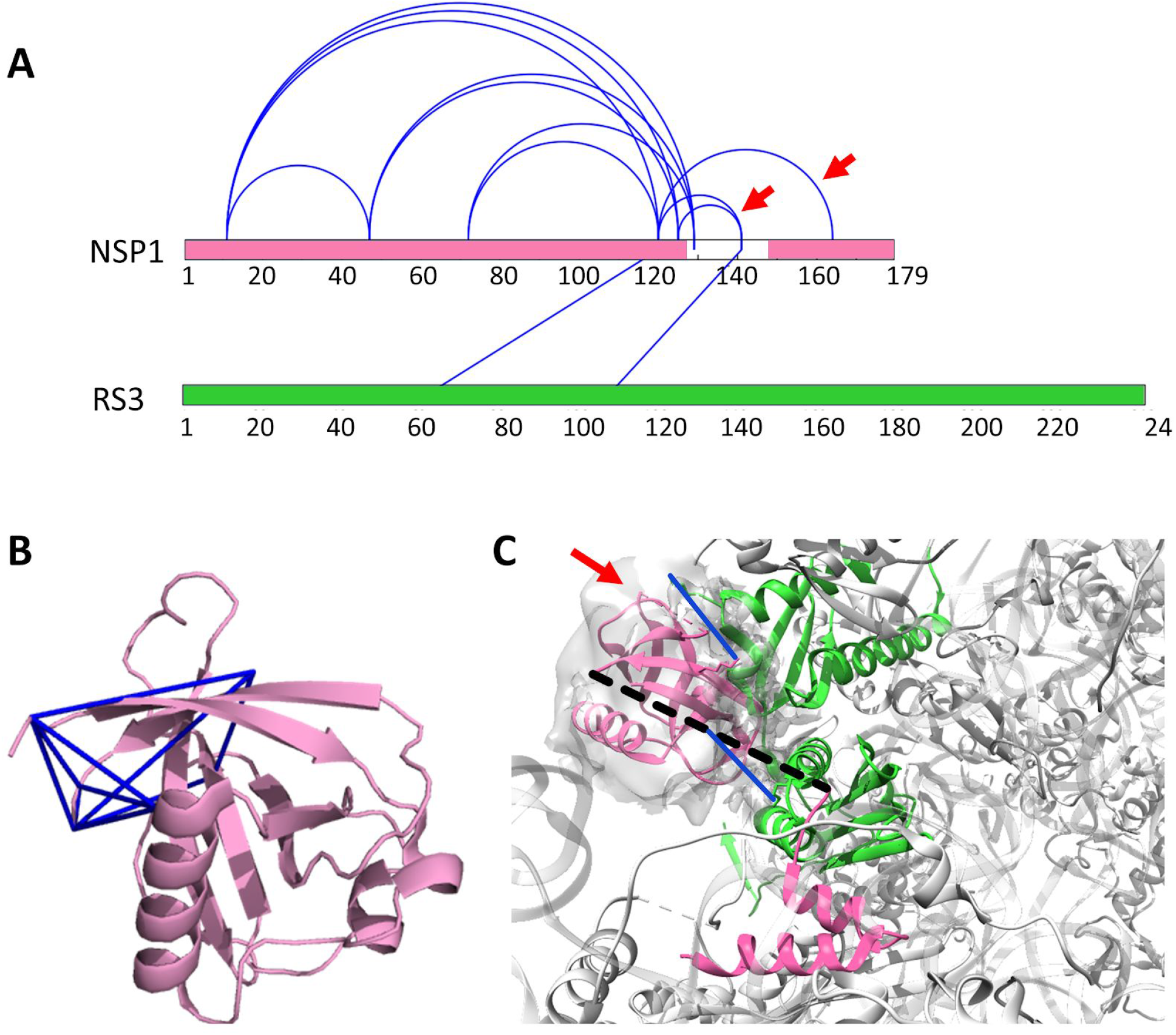
Nsp1 cross-links. **(A)** *In situ* cross-links identified within Nsp1, and between Nsp1 and ribosomal protein RS3. The two red arrows indicate cross-links that are not consistent with the structure of the C-terminal of Nsp1 bound to the ribosome. **(B)** *In situ* cross-links (blue lines) within Nsp1 mapped onto the crystal structure of the N-terminal domain (PDB 7k3n^45^). **(C)** Mapping of the Nsp1-RS3 crosslinks (blue) onto the cryo-EM structure of the C-terminal of Nsp1 (pink) bound to the 40S ribosomal subunit (gray) (PDB 6lzw^46^). RS3 is marked in green. The crystal structure of the N-terminal is docked into the unassigned density of the structure (red arrow). The unstructured linker between the N- and C-domains of Nsp1 is depicted by a dashed line. One of the cross-links involves a lysine within the unstructured linker.

Three *in situ* cross-links are not compatible with the structure of the C-terminal lodged deep within the mRNA entry tunnel (120 to 141; 125 to 141; 120 to 164). Because the bulky N-terminal cannot enter the mRNA entry tunnel, the occurrence of these cross-links implies an additional conformation of Nsp1 in which the C-terminal is interacting with the N-terminal. It is suggestive of an auto-inhibition mechanism for Nsp1 in which the N-terminal modulates the availability of the C-terminal towards the interaction with the ribosome.

## Discussion

The results establish the effectiveness of targeted *in situ* CLMS and integrative structure modeling to study a variety of proteins. The large number of identified cross-links allowed us to apply integrative structure modeling for domain assembly (Nsp2), oligomerization and domain assembly (N), and complex assembly (Nsp1-ribosome). Integration of domain level models generated by deep learning methods (AlphaFold2) with *in situ* CLMS data enabled *ab initio* structure modeling of the relatively long Nsp2 (638 amino acids).

Two factors contributed significantly to the identification yield of the *in situ* CLMS. The first is the use of the highly hydrophobic DSS reagent for cross-linking. The hydrophobicity improves the membrane permeability, thereby allowing to shorten the incubation time considerably. In our opinion, shortening the incubation times is crucial for maintaining the native cell state while minimizing the toxic effects of the cross-linker. The second factor is the utilization of the Strep tag technology that proved to be both highly effective for purification and highly compatible with CLMS.

In contrast to the success of *in situ* CLMS to identify dozens of intra-protein cross-links within the three SARS-CoV-2 proteins, the number of inter-protein cross-links was significantly lower. Cross-links between Nsp2 and Prohibitin were not identified, nor were cross-links between Nsp1 and other ribosomal proteins beside RS3. Three reasons may underlie these missing identifications. First, these proteins may be intended to interact strongly with other viral proteins that are not present without the context of a full viral infection. Second, the lack of ‘cross-linkable’ residues at the interacting interfaces of these particular systems. If this is the case, then the use of membrane-permeable reagents that employ other cross-linking chemistries (e.g. UV-induced, etc.) may be beneficial in future studies. Third, the sub-stoichiometric nature of the virus-host interactions reduces the signal of inter-protein cross-links. We note that in the opposite case, Wang, *et al*.^20–22^ targeted a fully stoichiometric complex (the proteasome) and identified inter-protein cross-links with high yield.

A general observation from this study is the large variation between *in situ* and *in vitro* experiments. For both Nsp2 (**Figure 2D**) and N (**Figure S7**) the overlap between the cross-link sets is partial, even though the underlying chemical reactivity is identical. One would expect the *in vitro* sets to fully contain the *in situ* sets, because the formers are not encumbered by issues of membrane-permeability. Yet, notably, a considerable number of cross-links are found only in the *in situ* sets. We interpret these results to indicate that *in situ* cross-linking probes protein states that are not occuring *in vitro*. These states may require certain cellular factors that are depleted upon cell lysis, thereby leading to denaturation, aggregation, or oligomer disassembly. Overall, the *in situ*/*in vitro* disparity highlights the importance of developing *in situ* techniques for the study of recalcitrant proteins. Accordingly, we believe that future *in situ* CLMS experiments in the context of a full viral infection would provide an even more informative picture on the functions of these proteins.

## Methods

### Cloning

The plasmid for expression of the Nucleocapsid protein is based on the pcDNA3.4 backbone (Thermo). cDNA of SARS-CoV2 was generated from a clinical RNA sample (Hadassah Medical Center, Clinical Virology laboratory, Prof. Dana Wolf), using QuantaBio qscript cDNA synthesis kit, followed by amplification of Nucleocapsid coding region by specific primers. The amplified PCR fragment of Nucleocapsid coding region was subsequently cloned using Gibson assembly reaction into pcDNA3.4 backbone modified to include C-terminal Strep Tag-II (IBA). Sequence of the cloned Nucleocapsid with C-terminal Strep tag was verified using Sanger sequencing. The sequence is identical to the canonical Nucleocapsid sequence (UniProt:P0DTC9). Plasmids for expression of Nsp1 and Nsp2 with a Strep-tag were kindly provided by the Krogan lab^48^. All plasmids were amplified under ampicillin selection in Top10 cells (Invitrogen) and purified by PureLink (Invitrogen).

### Cell culture and transfection

Human embryonic kidney cells 293 (HEK293, ATCC) were cultured (DMEM High glucose 10% FBS) at 37°C, 5% CO2, and high humidity. Three days prior to the transfection, the cells were plated in a 10 cm plate at an initial density of 2.25 × 10^6^ cells/plate. Expression plasmids (Nsp1/Nsp2/N) and PEI (Polysciences, 260008-5) were separately diluted in Opti-MEM1 (Gibco 31985-047) and mixtures were incubated at room temperature for 25 minutes to allow polyplex formation prior to its addition to the cell culture. The mixture was added dropwise onto the cells. The plates were washed and fed with fresh medium 24 hours post-transfection. The cells were dissociated 40 hours after transfection by application of Dulbecco’s Phosphate Buffered Saline without calcium and magnesium (D-PBS) supplemented with 10 mM EDTA for 5 minutes at 37°C. The cells were pelleted and transferred into a 1.7 ml tube. Pelleting of intact cells was always carried out by centrifugation at 200 g for three minutes at either room temperature (for buffers at 37°C) or 4°C (for ice-cold buffers).

### *In situ* cross-linking

Stock solution of 250 mM DSS (disuccinimidyl suberate) in DMSO was freshly prepared on the day of the experiment. Stock solution of formalin (37% formaldehyde, 10% methanol) was prepared in the month of the experiment. The dissociation buffer was removed and replaced by the cross-linking buffer, which was warm D-PBS supplemented with either 0.3% formaldehyde or 10 mM DSS. Note, the mixing of DSS with D-PBS resulted immediately in a cloudy solution because the solubility of DSS in aqueous buffer is far less than 10 mM. The rationale for the value of 10 mM was to ensure that the DSS reservoir in the buffer is not depleted during the cross-linking incubation. The cells were incubated with the cross-linker for 20 minutes at 37°C under constant gentle agitation to ensure that a cell pellet is not forming. The cells were pelleted and the cross-linking buffer was replaced with ice-cold quenching buffer (50 mM Tris-HCl, pH 7.5, 150 mM NaCl, 1 mM EDTA) to inactivate excess cross-linker around the intact cells. Quenching proceeded at 4°C for 10 minutes under gentle agitation. The cells were pelleted and the quenching buffer removed.

### Affinity purification

The cells were resuspended in 600 μl of lysis buffer (50 mM Tris-HCl, pH 7.5, 150 mM NaCl, 1 mM EDTA, 0.5% Triton X-100, 1% Protease inhibitor cocktail (Sigma P8340)). Lysis was performed by sonication on ice (1 minute total, 50% amplitude, cycle: 5 second on 25 second off, Qsonica Q125 with 1/8’’ probe) followed by centrifugation at 13,000 g, 4°C for 10 minutes to pellet debris. The supernatant was incubated at 4°C for 4 hours with 10 μl of StrepTactin resin (Sepharose High Performance, Cytiva). The lysate was removed, and the resin was manually washed three times with 1 ml of wash buffer (50 mM Tris-HCl, pH 7.5, 150 mM NaCl, 1 mM EDTA). Throughout the washing steps, the resin was brought to the bottom of the tube by centrifugation at 200 g for 45 seconds. The overall duration of the washing step was 6 minutes. To elute the protein, the beads were covered with wash buffer supplemented with 10 mM biotin for 30 minutes with occasional gentle mixing. The supernatant was collected for mass spectrometry.

### *In vitro* cross-linking

For *in vitro* cross-linking, the dissociated cells were immediately resuspended in the lysis buffer and lysed as described above. The affinity purification was modified to use HEPES Wash Buffer (50 mM HEPES-HCl, pH 8.0, 150 mM NaCl, 1 mM EDTA) and HEPES Elution Buffer (HEPES Wash Buffer with 10 mM biotin). For BS3 (bis(sulfosuccinimidyl)suberate) cross-linking, the eluted protein was incubated with 1 mM BS3 at 30°C for 30 minutes, quenched with 20 mM ammonium bicarbonate for 10 minutes, and then prepared for mass spectrometry. For DMTMM cross-linking, we did not elute the protein from the StrepTactin resin, because of concerns for the possible interaction of biotin with DMTMM. Instead, we performed on-bead cross-linking by incubating the resin with 100 μl HEPES Wash Buffer supplemented with 10 mM DMTMM at 30°C for 30 minutes. The buffer with the cross-linker was removed, and on-bead digestion was employed to prepare the sample for mass spectrometry.

### Preparation of samples for mass spectrometry

The proteins were precipitated in 1 ml of acetone (−80 °C) for one hour, followed by centrifugation at 13,000 g. The pellet was resuspended in 20 μl of 8 M urea with 10 mM DTT. After 30 minutes, iodoacetamide was added to a final concentration of 25 mM and the alkylation reaction proceeded for 30 minutes. The urea was diluted by adding 200 μl of Digestion Buffer (25 mM TRIS, pH=8.0; 10% Acetonitrile). We added 0.5 μg of trypsin (Promega) to the diluted urea and digested the protein overnight at 37°C under agitation. Following digestion, the peptides were desalted on C18 stage-tips and eluted by 55% acetonitrile. The eluted peptides were dried in a SpeedVac, reconstituted in 0.1% formic acid, and measured in the mass spectrometer.

Based on UV absorbance measurements, the final amounts per experiment of peptides in the tryptic digest were 20 μg, 2 μg, and 200 ng for N, Nsp2, and Nsp1, respectively. Despite the low amount, we attempted to enrich for cross-linked peptides in the case of Nsp2. To that end, peptides from two different purifications were enriched by either strong cation exchange chromatography^49^ or size exclusion chromatography^50^. The enrichments led to the identification of three new cross-links (out of 53) over the standard mass spectrometric analysis of the full digest. We conclude that the common enrichment techniques are not effective for protein samples of such low amounts.

### Mass spectrometry

The samples were analyzed by a 120 minute 0-40% acetonitrile gradient on a liquid chromatography system coupled to a Q-Exactive HF mass spectrometer. The analytical column was an EasySpray 25 cm heated to 40°C. The method parameters of the run were: Data-Dependent Acquisition; Full MS resolution 70,000; MS1 AGC target 1e6; MS1 Maximum IT 200 ms; Scan range 450 to 1800; dd-MS/MS resolution 35,000; MS/MS AGC target 2e5; MS2 Maximum IT 600 ms; Loop count Top 12; Isolation window 1.1; Fixed first mass 130; MS2 Minimum AGC target 800; Peptide match - off; Exclude isotope - on; Dynamic exclusion 45 seconds. Each cross-linked sample was measured twice in two different HCD energies (NCE): 26, and stepped 25, 30, and 35. All cross-linked samples were measured with the following charge exclusion: unassigned,1,2,3,8,>8. Proteomics samples were measured with the following charge exclusion: unassigned,1,8,>8.

### Proteomics analysis of interacting proteins

Human proteins that interact with the bait proteins were identified by comparing the protein content between purifications from transfected and untransfected cells. Label-free quantification (LFQ) was performed with MaxQuant 1.5^51^ using the default parameters. The sequence database comprised all human proteins (downloaded from UniProt) augmented with the sequences of three SARS-CoV-2 proteins: Nsp1, Nsp2, and Nucleocapsid protein. The ‘proteinGroups.txt’ output file was loaded to Perseus^52^. Reverse proteins and contaminations were filtered out, the data were transformed to logarithmic scale, and samples grouped according to replicates. Only proteins identified by more than two peptides were considered. For missing LFQ intensities, values were imputed from a normal distribution. The confidence curves were determined by a two-sample test with a permutation-based FDR of 0.5% and ‘s0’ (minimal fold change) value of 2. The Volcano plots present the proteins in the “t-test difference” vs. “-Log t-test p-value” coordinate system.

### Identification of cross-links

The RAW data files from the mass spectrometer were converted to MGF format by Proteome Discoverer (Thermo). The identification of DSS cross-links used a search application^53^ that exhaustively enumerates all the possible peptide pairs. The search parameters were as follows: Sequence database – Proteins above the confidence line in the volcano plots (Figure S1); Protease – trypsin, allowing up to three miscleavage sites; Fixed modification of cysteine by iodoacetamide; Variable modifications: methionine oxidation, lysine with hydrolyzed mono-link; Cross-linking must occur between two lysine residues; Cross-linker is never cleaved; MS/MS fragments to consider: b-ions, y-ions; MS1 tolerance – 6 ppm; MS2 tolerance – 8 ppm; Cross-linker mass – one of three possible masses: 138.0681, 138.0681 + 1.00335, and 138.0681 + 2.0067. The three masses address the occasional incorrect assignment of the mono-isotopic mass by the mass spectrometer^54^.

A cross-link was identified as a match between a measured MS/MS event and a peptide pair if it fulfilled four conditions: 1) The mass of the precursor ion is within the MS1 tolerance of the theoretical mass of the linked peptide pair (with either of the three possible cross-link masses); 2) At least four MS/MS fragments were identified within the MS2 tolerance on each peptide; 3) The fragmentation score of the cross-link (defined as the number of all matching MS/MS fragments divided by the combined length of the two peptides) is above a threshold determined by the FDR analysis (see ahead); 4) The fragmentation score is at least 15% better than the score of the next best peptide pair or linear peptide.

### Estimation of the false detection rate (FDR)

The FDR was estimated from decoy-based analysis. To that end, the cross-link identification analysis was repeated 20 times with an erroneous cross-linker mass of 138.0681*N/138 Da, where N=160, 161, 162, … 179. This led to bogus identifications with fragmentation scores that were generally much lower than the scores obtained with the correct cross-linker mass (see histograms in **Figure S3**). For the identification of true cross-links, we set the threshold on the fragmentation score according to the desired FDR value. For example, a threshold of 0.65 on the fragmentation score of the Nsp2 data set gave 53 cross-links above the threshold in the true analysis, and a median of 1 cross-link in a typical decoy run (**Figure S2C**). We therefore estimate the corresponding FDR to be about 1 in 53, or ∼2%. The final thresholds used: Nsp2 Primary Set – 0.65, Nsp2 Secondary Set – 0.9, Nsp2 *in vitro* Set – 0.65, N – 0.7, Nsp1 – 0.65.

### Domain assembly via pairwise docking

The input to the domain assembly problem consists of a set of structural models and a list of cross-links. The goal is to predict an assembly with good complementarity between the domains and consistency with the input cross-links. We use CombDock^33^, a combinatorial docking algorithm, which was modified to support cross-linking data^35^. First, pairwise docking is applied on each pair of input structures to generate a set of docked configurations (**Step 1**). Second, combinatorial optimization is used to combine different subsets of the configurations from pairwise docking to generate clash-free complex models consistent with the cross-links and chain connectivity (**Step 2**). A cross-link is considered satisfied if the distance between the Cα atoms of the cross-linked residues is below a specified threshold. Here we used a threshold of 25 Å and 20 Å for DSS and formaldehyde cross-links, respectively.

### Step 1. All-pairs docking

We used PatchDock to generate pairs of docked configurations^34^. PatchDock employs an efficient rigid docking algorithm that maximizes geometric shape complementarity. Protein flexibility is accounted for by a geometric shape complementarity scoring function, which allows a small amount of steric clashes at the interface. Only domain pairs with at least one cross-link between them were structurally docked at this stage. The PatchDock scoring function was augmented by restraints derived from the cross-linking data. Each docking configuration is represented by a transformation (three rotational and three translational parameters) and we keep the K=1,000 best scoring transformations for each pair of domains.

### 2. Combinatorial optimization

Two basic principles are used by the algorithm: a hierarchical construction of the assembly and a greedy selection of subcomplexes. The input comprises the pair-wise docking of Step 1 (subcomplexes of size 2). At each step, the algorithm generates subcomplexes with n subunits by connecting two subcomplexes of smaller size. Only valid subcomplexes are retained at each step. Valid subcomplexes do not contain steric clashes and satisfy distance constraints (chain connectivity) and restraints (cross-links). Searching the entire space is impractical, even for relatively small K (number of models per pair) and N (number of domains), due to computer speed and memory limitations. Therefore, the algorithm performs a greedy selection of subcomplexes by keeping only the D=1,000 best-scoring models at each step. The final models are clustered using RMSD clustering with a cutoff of 4 Å. The final best scoring models are selected based on the cross-links satisfaction and cluster size. The model precision is calculated as the average Cα RMSD between the best-scoring models.

Pairwise docking was applied to dock the RNA binding domain of N (PDB 7act) to the dimerization domain (PDB 6zco). Combinatorial optimization was performed for assembly of Nsp2 domains.

### Molecular dynamics (MD) simulations

MD simulations were performed on the dimerization domain of the N protein model with docked RNA using GROMACS 2020 software^55^ and the PARMBSC1 force field^56^. Any steric clashes between the docked RNA and protein were resolved by removing nucleotide bases, leaving a poly-U hexameric RNA fragment. Then the protein-RNA complex was solvated with a simple point charge water model with a self-energy polarization correction term (SPC/E)^57^, and the system charge was neutralized with the addition of Cl^-^ ions. In order to ensure appropriate initial geometry, a steepest-descent energy minimization MD run was allowed to run until convergence at Fmax < 1000 kJ/(mol · nm). Position restraints with k = 1000 kJ/(mol · nm^2^) are then applied to the protein and RNA heavy atoms to allow the water and ions to equilibrate around the protein in two-steps. The first equilibration step is conducted at a constant number of atoms, volume, and temperature (NVT) of 300°K. The second equilibration step is conducted at a constant number of atoms, pressure of 1 bar, and temperature of 300°K (NPT). These equilibrations have a time step of 2 fs and last for 100 ps. Once the system is equilibrated at 300°K and 1 bar, a production simulation of 100 nanoseconds with a time step of 2 fs provides 10,000 MD simulation frames at intervals of 10ps.

### Data Availability

The mass spectrometry data have been deposited to the ProteomeXchange Consortium via the PRIDE^58^ partner repository.

The cross-link information is compiled with the dataset identifier PXD023487. Reviewers can access the data with the following:

Username

Password: xKhhA75i

The proteomics information is compiled with the dataset identifier PXD023542. Reviewers can access the data with the following:

Username

Password: Yu2juXAt

## Supporting information

Model of N protein

Model of Nsp2

## Supplementary Figures

**Supplementary Figure 1.**
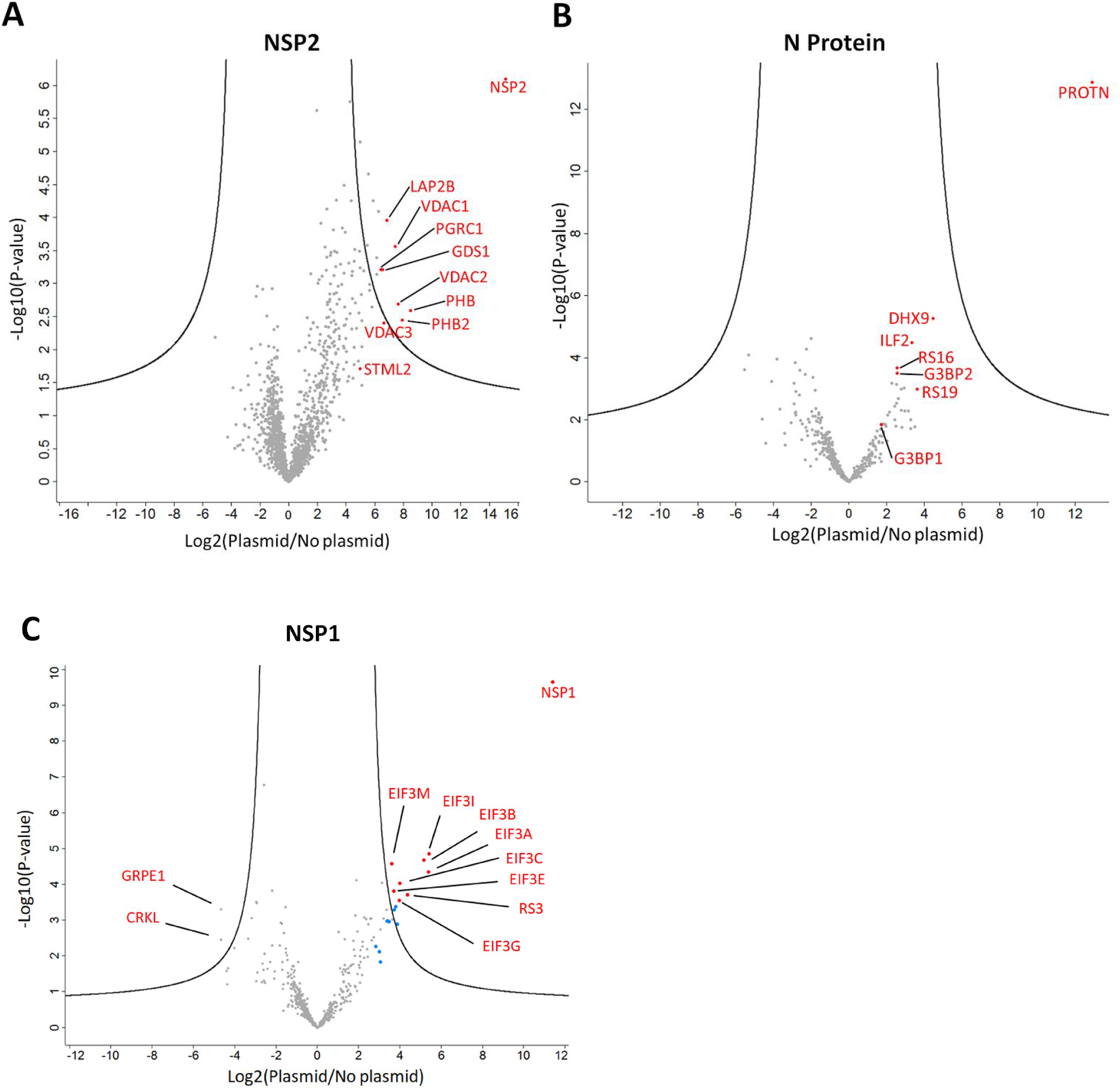
Volcano plots quantifying the protein compositions of purifications from transfected vs. untransfected cells. FDR thresholds of 0.5% are marked by the black curves. **(A)** Nsp2 transfection. Components of the Prohibitin complex (PHB, VDAC, and STML2) co-elute in significant amounts. **(B)** N protein transfection. The N protein does not appear to have a specific human protein interactor **(C)** Nsp1 transfection. Components of the 40S ribosome co-eluted with Nsp1. Interestingly, the CCT chaperonin was abundant in the Nsp1 elution (blue dots mark the CCT

**Supplementary Figure 2.**
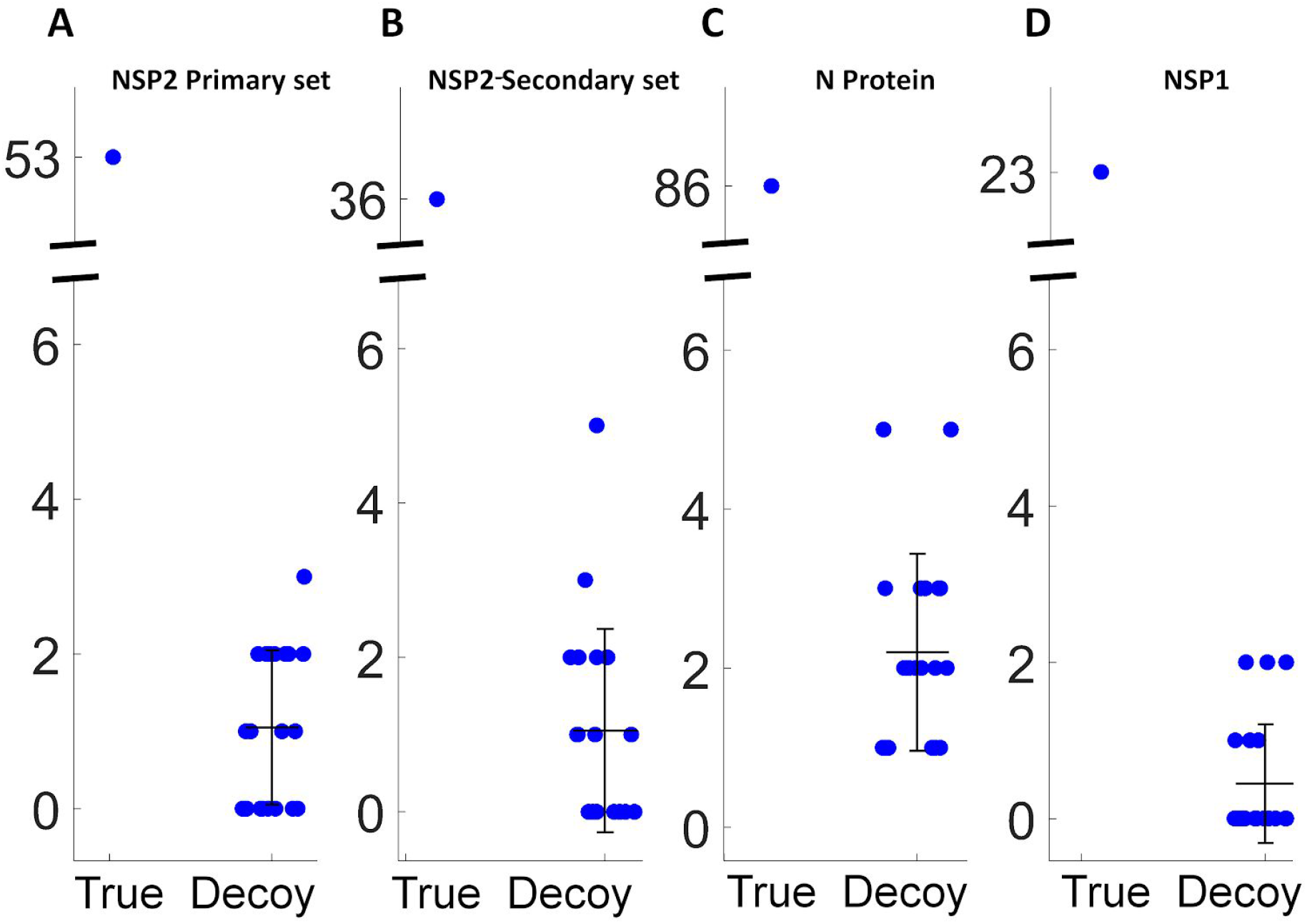
Estimation of the false-detection rate (FDR) for the DSS cross-link sets. The identification analysis was repeated 20 times with an erroneous cross-linker mass that differed from the true mass (138.0681 Da) by 20 to 39 Da. Each blue dot represents the number of identified cross-links from either a true or decoy analysis. The mean and standard deviation of each decoy set is shown. The FDR for each set was the ratio of the median value of the decoys to the number of true identifications.

**Supplementary Figure 3.**
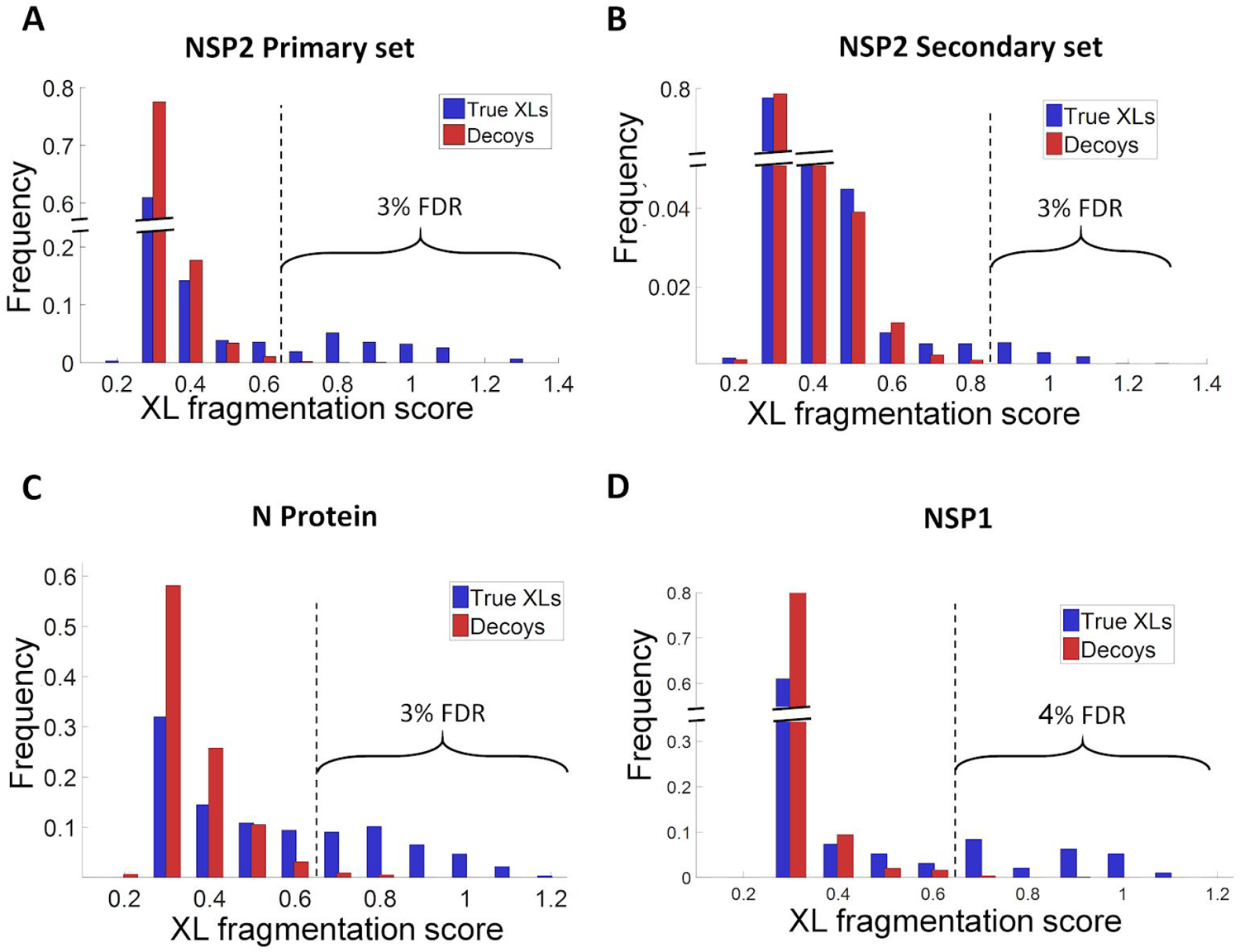
Distributions of the XL fragmentation score (number of MS/MS fragments divided by the total length of the peptides). For each cross-link set, we report only the cross-links above a certain score threshold that is marked by a dashed line. **(A)** For the primary set of Nsp2 the threshold is 0.65, which sets an FDR of ∼3%. **(B)** For the secondary set of Nsp2 the threshold is 0.9 **(C)** For the N protein the threshold is 0.7 **(D)** For Nsp1 the threshold is 0.65

**Supplementary Figure 4.**
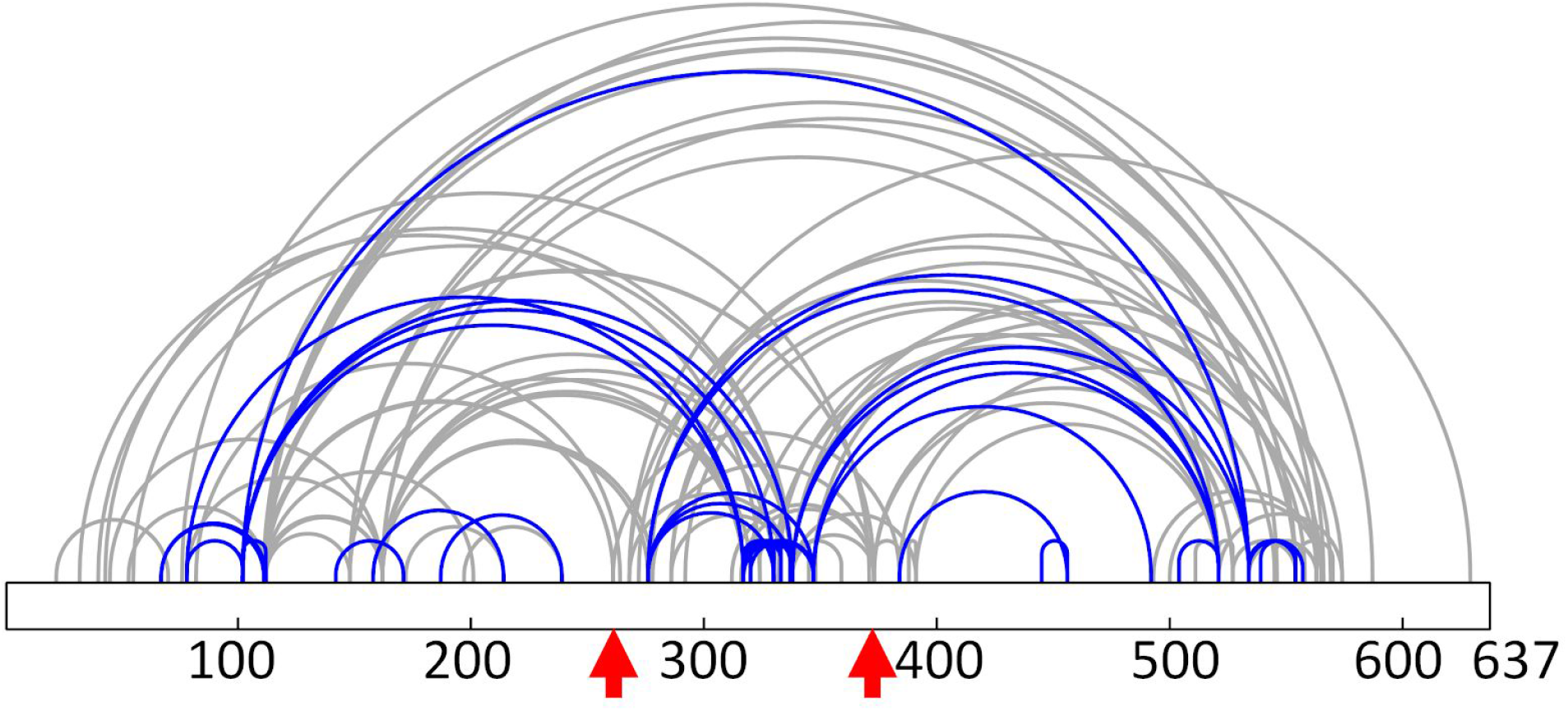
Comparison of cross-links identified within Nsp2 from *in situ* CLMS using DSS (blue) and *in vitro* CLMS using DMTMM (grey). Similarly to BS3 (**Figure 2C**), *in vitro* experiments with DMTMM caused some lysine residues to link with the entire protein sequence (red arrows).

**Supplementary Figure 5.**
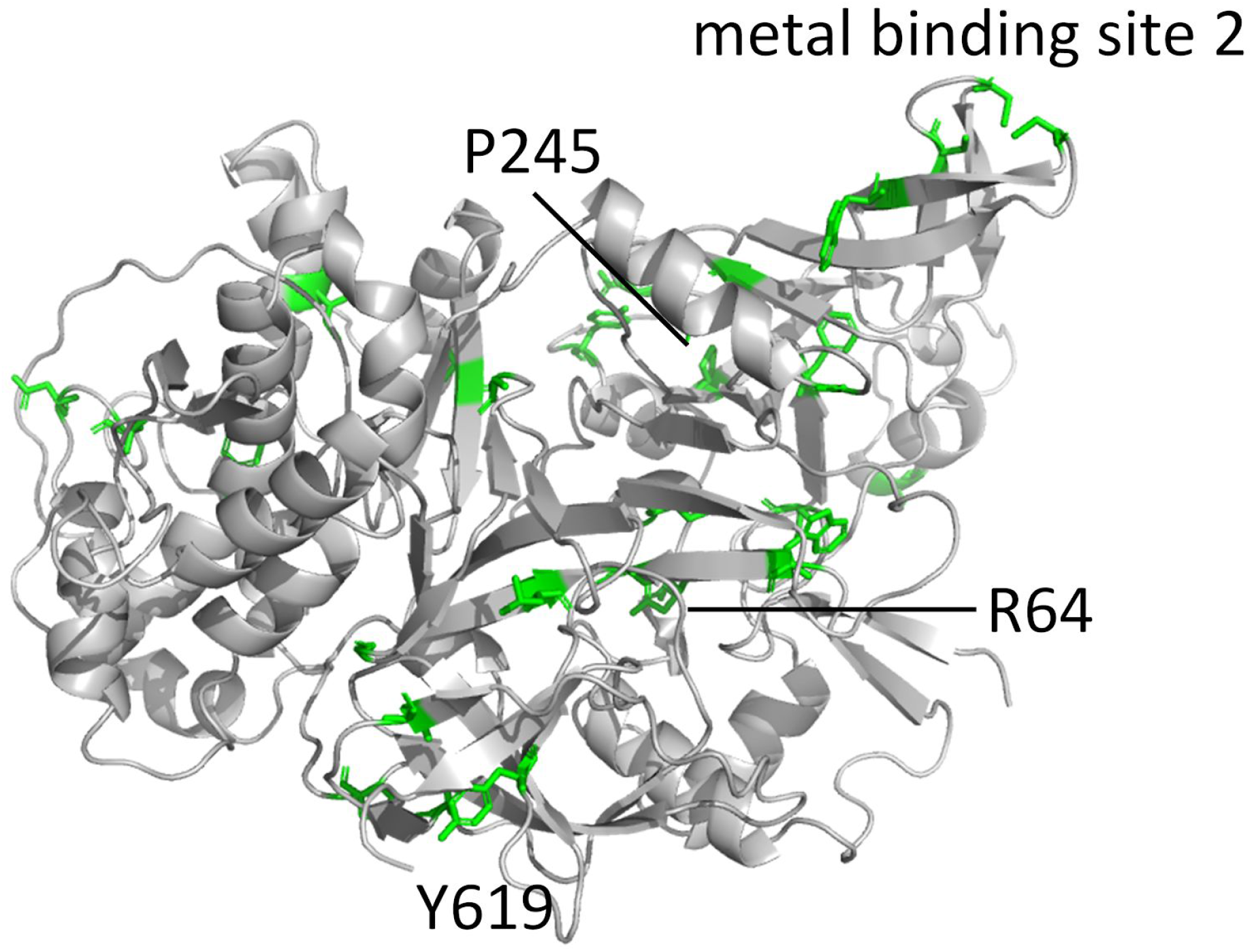
Mapping of highly conserved residues of Nsp2 (green) across all human coronaviruses onto the full-length integrative model. Clusters of conservation are apparent around Pro245 and Tyr619.

**Supplementary Figure 6.**
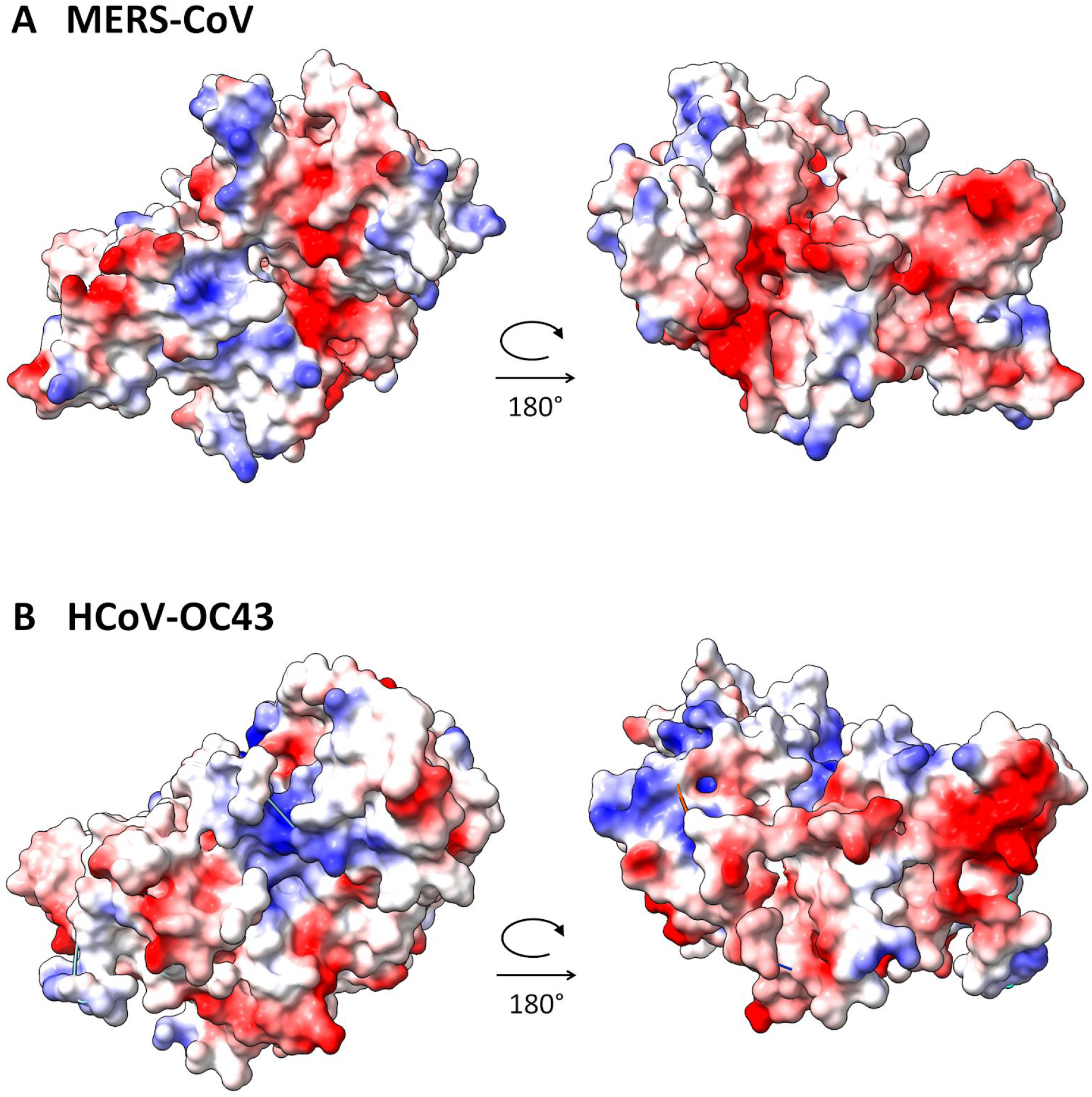
Electrostatic surfaces of homology models of Nsp2 sequences of MERS and OC43 that are based on our integrative model as template. The large acidic patch (right views) is conserved across human coronaviruses.

**Supplementary Figure 7.**
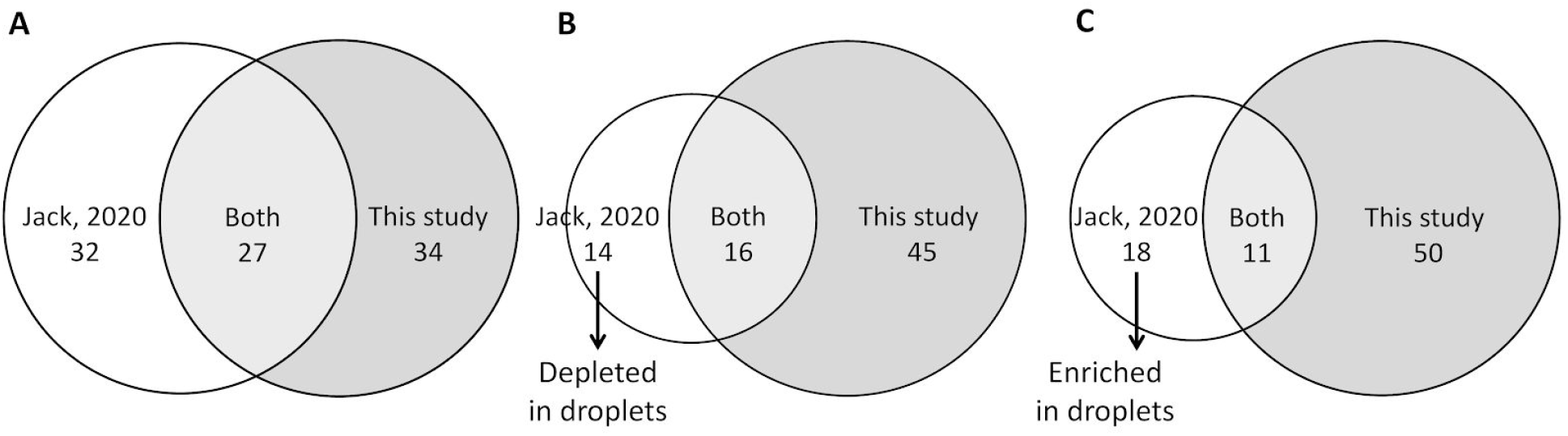
Overlap of N protein cross-link sets obtained under *in vitro* conditions (Jack *et al*.^41^, white) or *in situ* (this study, grey). Jack *et al*.^41^ cross-linked the N protein under two conditions – soluble N protein (at high salt concentration) and phase separated N protein in droplets (at low salt concentration). They report **(A)** 59 cross-links overall, **(B)** 30 cross-links that are depleted in the phase separated condition, and **(C)** 29 cross-links that are enriched in the phase separated condition. The *in situ* cross-links seem to originate from both conditions.

## References

1. Bojkova, D. et al.. Proteomics of SARS-CoV-2-infected host cells reveals therapy targets. Nature 583, 469–472 (2020).

2. Finkel, Y. et al.. The coding capacity of SARS-CoV-2. Nature 589, 125–130 (2021).

3. Kim, D. et al.. The Architecture of SARS-CoV-2 Transcriptome. Cell 181, 914–921.e10 (2020).

4. Sedova, M., Jaroszewski, L., Alisoltani, A. & Godzik, A. Coronavirus3D: 3D structural visualization of COVID-19 genomic divergence. Bioinformatics 36, 4360–4362 (2020).

5. Scudellari, M. The sprint to solve coronavirus protein structures - and disarm them with drugs. Nature 581, 252–255 (2020).

6. Bruce, J. E. In vivo protein complex topologies: sights through a cross-linking lens. Proteomics 12, 1565–1575 (2012).

7. Kaake, R. M. et al.. A new in vivo cross-linking mass spectrometry platform to define protein-protein interactions in living cells. Mol. Cell. Proteomics 13, 3533–3543 (2014).

8. Leitner, A., Faini, M., Stengel, F. & Aebersold, R. Crosslinking and Mass Spectrometry: An Integrated Technology to Understand the Structure and Function of Molecular Machines. Trends Biochem. Sci. 41, 20–32 (2016).

9. Sinz, A. Cross-Linking/Mass Spectrometry for Studying Protein Structures and Protein-Protein Interactions: Where Are We Now and Where Should We Go from Here? Angew. Chem. Int. Ed Engl. 57, 6390–6396 (2018).

10. Schneider, M., Belsom, A. & Rappsilber, J. Protein Tertiary Structure by Crosslinking/Mass Spectrometry. Trends Biochem. Sci. 43, 157–169 (2018).

11. Yu, C. & Huang, L. Cross-Linking Mass Spectrometry: An Emerging Technology for Interactomics and Structural Biology. Anal. Chem. 90, 144–165 (2018).

12. Shi, Y. et al.. A strategy for dissecting the architectures of native macromolecular assemblies. Nat. Methods 12, 1135–1138 (2015).

13. Schweppe, D. K. et al.. Mitochondrial protein interactome elucidated by chemical cross-linking mass spectrometry. Proc. Natl. Acad. Sci. U. S. A. 114, 1732–1737 (2017).

14. Fasci, D., van Ingen, H., Scheltema, R. A. & Heck, A. J. R. Histone Interaction Landscapes Visualized by Crosslinking Mass Spectrometry in Intact Cell Nuclei. Mol. Cell. Proteomics 17, 2018–2033 (2018).

15. Liu, F., Lössl, P., Rabbitts, B. M., Balaban, R. S. & Heck, A. J. R. The interactome of intact mitochondria by cross-linking mass spectrometry provides evidence for coexisting respiratory supercomplexes. Mol. Cell. Proteomics 17, 216–232 (2018).

16. Ryl, P. S. J. et al.. In Situ Structural Restraints from Cross-Linking Mass Spectrometry in Human Mitochondria. J. Proteome Res. 19, 327–336 (2020).

17. Gonzalez-Lozano, M. A. et al.. Stitching the synapse: Cross-linking mass spectrometry into resolving synaptic protein interactions. Sci Adv 6, eaax5783 (2020).

18. Navare, A. T. et al.. Probing the protein interaction network of Pseudomonas aeruginosa cells by chemical cross-linking mass spectrometry. Structure 23, 762–773 (2015).

19. O’Reilly, F. J. et al.. In-cell architecture of an actively transcribing-translating expressome. Science 369, 554–557 (2020).

20. Wang, X. et al.. Molecular Details Underlying Dynamic Structures and Regulation of the Human 26S Proteasome. Mol. Cell. Proteomics 16, 840–854 (2017).

21. Wang, X. et al.. The proteasome-interacting Ecm29 protein disassembles the 26S proteasome in response to oxidative stress. J. Biol. Chem. 292, 16310–16320 (2017).

22. Tayri-Wilk, T. et al.. Mass spectrometry reveals the chemistry of formaldehyde cross-linking in structured proteins. Nat. Commun. 11, 3128 (2020).

23. Chavez, J. D. et al.. Chemical Crosslinking Mass Spectrometry Analysis of Protein Conformations and Supercomplexes in Heart Tissue. Cell Syst 6, 136–141.e5 (2018).

24. Rout, M. P. & Sali, A. Principles for Integrative Structural Biology Studies. Cell 177, 1384–1403 (2019).

25. Koukos, P. I. & Bonvin, A. M. J. J. Integrative Modelling of Biomolecular Complexes. J. Mol. Biol. 432, 2861–2881 (2020).

26. Narayanan, K. et al.. Severe acute respiratory syndrome coronavirus nsp1 suppresses host gene expression, including that of type I interferon, in infected cells. J. Virol. 82, 4471–4479 (2008).

27. Graham, R. L., Sims, A. C., Brockway, S. M., Baric, R. S. & Denison, M. R. The nsp2 replicase proteins of murine hepatitis virus and severe acute respiratory syndrome coronavirus are dispensable for viral replication. J. Virol. 79, 13399–13411 (2005).

28. Hagemeijer, M. C. et al.. Dynamics of coronavirus replication-transcription complexes. J. Virol. 84, 2134–2149 (2010).

29. Jones, D. T. Protein secondary structure prediction based on position-specific scoring matrices. J. Mol. Biol. 292, 195–202 (1999).

30. Cornillez-Ty, C. T., Liao, L., Yates, J. R., 3rd, Kuhn, P. & Buchmeier, M. J. Severe acute respiratory syndrome coronavirus nonstructural protein 2 interacts with a host protein complex involved in mitochondrial biogenesis and intracellular signaling. J. Virol. 83, 10314–10318 (2009).

31. Callaway, E. ‘It will change everything’: DeepMind’s AI makes gigantic leap in solving protein structures. Nature vol. 588 203–204 (2020).

32. Li, Y., Ren, Z., Bao, Z., Ming, Z. & Li, X. Expression, crystallization and preliminary crystallographic study of the C-terminal half of nsp2 from SARS coronavirus. Acta Crystallogr. Sect. F Struct. Biol. Cryst. Commun. 67, 790–793 (2011).

33. Inbar, Y., Benyamini, H., Nussinov, R. & Wolfson, H. J. Combinatorial docking approach for structure prediction of large proteins and multi-molecular assemblies. Phys. Biol. 2, S156–65 (2005).

34. Schneidman-Duhovny, D., Inbar, Y., Nussinov, R. & Wolfson, H. J. Geometry-based flexible and symmetric protein docking. Proteins 60, 224–231 (2005).

35. Schneidman-Duhovny, D. & Wolfson, H. J. Modeling of Multimolecular Complexes. Methods Mol. Biol. 2112, 163–174 (2020).

36. Holm, L. Using Dali for Protein Structure Comparison. Methods Mol. Biol. 2112, 29–42 (2020).

37. Dinesh, D. C. et al.. Structural basis of RNA recognition by the SARS-CoV-2 nucleocapsid phosphoprotein. PLoS Pathog. 16, e1009100 (2020).

38. Zinzula, L. et al.. High-resolution structure and biophysical characterization of the nucleocapsid phosphoprotein dimerization domain from the Covid-19 severe acute respiratory syndrome coronavirus 2. Biochem. Biophys. Res. Commun. (2020) doi:10.1016/j.bbrc.2020.09.131.

39. Lu, S. et al.. The SARS-CoV-2 nucleocapsid phosphoprotein forms mutually exclusive condensates with RNA and the membrane-associated M protein. Nat. Commun. 12, 502 (2021).

40. Carlson, C. R. et al.. Phosphoregulation of Phase Separation by the SARS-CoV-2 N Protein Suggests a Biophysical Basis for its Dual Functions. Mol. Cell 80, 1092–1103.e4 (2020).

41. Jack, A. et al.. SARS CoV-2 nucleocapsid protein forms condensates with viral genomic RNA. bioRxiv (2020) doi:10.1101/2020.09.14.295824.

42. Yao, H. et al.. Molecular Architecture of the SARS-CoV-2 Virus. Cell 183, 730–738.e13 (2020).

43. Ye, Q., West, A. M. V., Silletti, S. & Corbett, K. D. Architecture and self-assembly of the SARS-CoV-2 nucleocapsid protein. Protein Sci. 29, 1890–1901 (2020).

44. Luger, K., Mäder, A. W., Richmond, R. K., Sargent, D. F. & Richmond, T. J. Crystal structure of the nucleosome core particle at 2.8 A resolution. Nature 389, 251–260 (1997).

45. Semper, C., Watanabe, N. & Savchenko, A. Structural characterization of nonstructural protein 1 from SARS-CoV-2. iScience 24, 101903 (2021).

46. Thoms, M. et al.. Structural basis for translational shutdown and immune evasion by the Nsp1 protein of SARS-CoV-2. Science 369, 1249–1255 (2020).

47. Schubert, K. et al.. SARS-CoV-2 Nsp1 binds the ribosomal mRNA channel to inhibit translation. Nat. Struct. Mol. Biol. 27, 959–966 (2020).

48. Gordon, D. E. et al.. A SARS-CoV-2 protein interaction map reveals targets for drug repurposing. Nature 583, 459–468 (2020).

49. Klykov, O. et al.. Efficient and robust proteome-wide approaches for cross-linking mass spectrometry. Nat. Protoc. 13, 2964–2990 (2018).

50. Leitner, A. et al.. Expanding the chemical cross-linking toolbox by the use of multiple proteases and enrichment by size exclusion chromatography. Mol. Cell. Proteomics 11, M111.014126 (2012).

51. Cox, J. & Mann, M. MaxQuant enables high peptide identification rates, individualized p.p.b.-range mass accuracies and proteome-wide protein quantification. Nat. Biotechnol. 26, 1367–1372 (2008).

52. Tyanova, S. et al.. The Perseus computational platform for comprehensive analysis of (prote)omics data. Nat. Methods 13, 731–740 (2016).

53. Kalisman, N., Adams, C. M. & Levitt, M. Subunit order of eukaryotic TRiC/CCT chaperonin by cross-linking, mass spectrometry, and combinatorial homology modeling. Proc. Natl. Acad. Sci. U. S. A. 109, 2884–2889 (2012).

54. Lenz, S., Giese, S. H., Fischer, L. & Rappsilber, J. In-Search Assignment of Monoisotopic Peaks Improves the Identification of Cross-Linked Peptides. J. Proteome Res. 17, 3923–3931 (2018).

55. Pronk, S. et al.. GROMACS 4.5: a high-throughput and highly parallel open source molecular simulation toolkit. Bioinformatics 29, 845–854 (2013).

56. Ivani, I. et al.. Parmbsc1: a refined force field for DNA simulations. Nat. Methods 13, 55–58 (2016).

57. Berendsen, H. J. C., Grigera, J. R. & Straatsma, T. P. The missing term in effective pair potentials. The Journal of Physical Chemistry vol. 91 6269–6271 (1987).

58. Perez-Riverol, Y. et al.. The PRIDE database and related tools and resources in 2019: improving support for quantification data. Nucleic Acids Res. 47, D442–D450 (2019).

