## Supplementary material for "Targeted *in situ* cross-linking mass spectrometry and integrative modeling reveal the architectures of Nsp1, Nsp2, and Nucleocapsid proteins from SARS-CoV-2": Figure S8

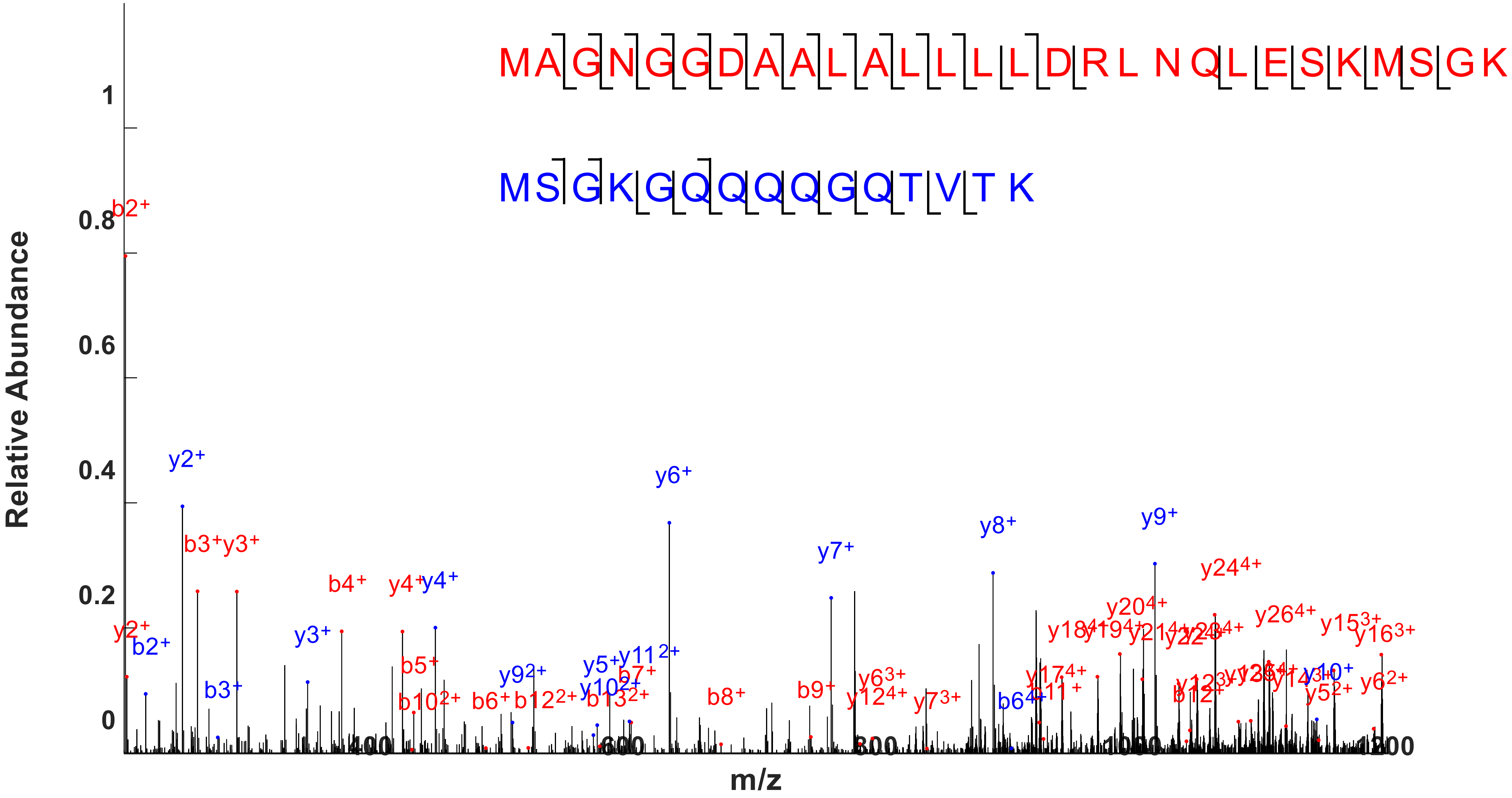


Protein N dimer cross link (1) K233 – K237.

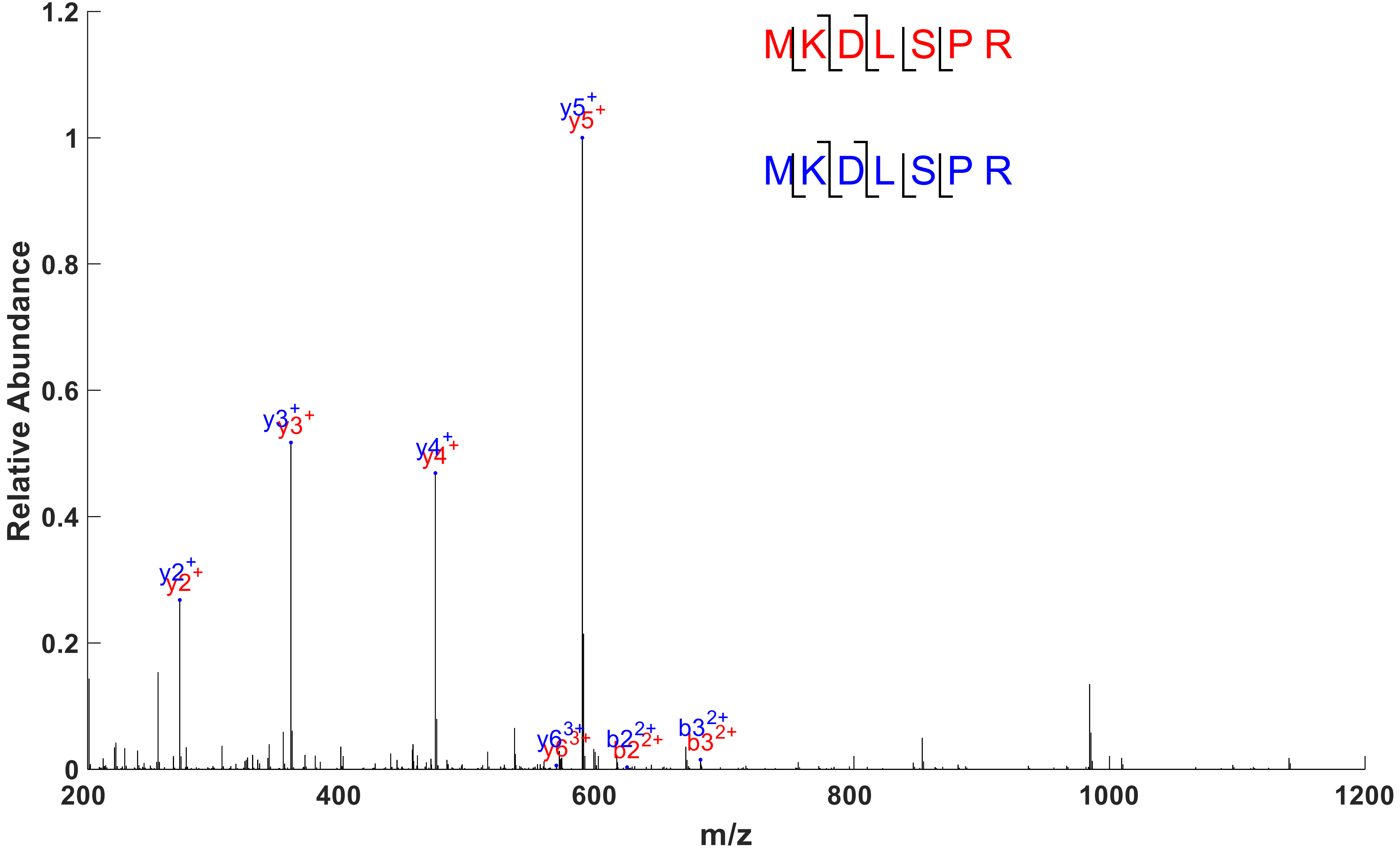


Protein N dimer cross link (2) K102 – K102.

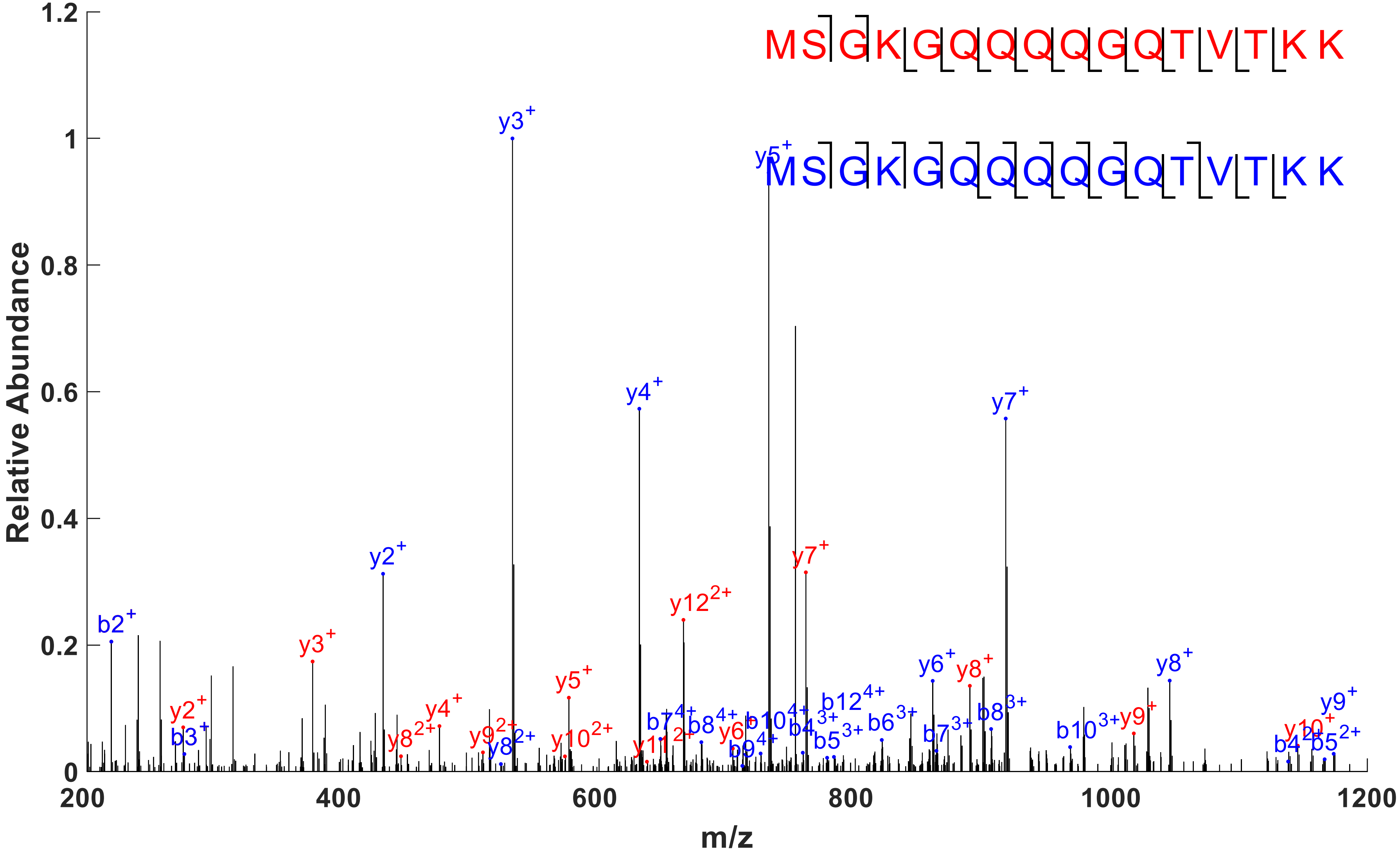


Protein N dimer cross link (3) K237 – K237.

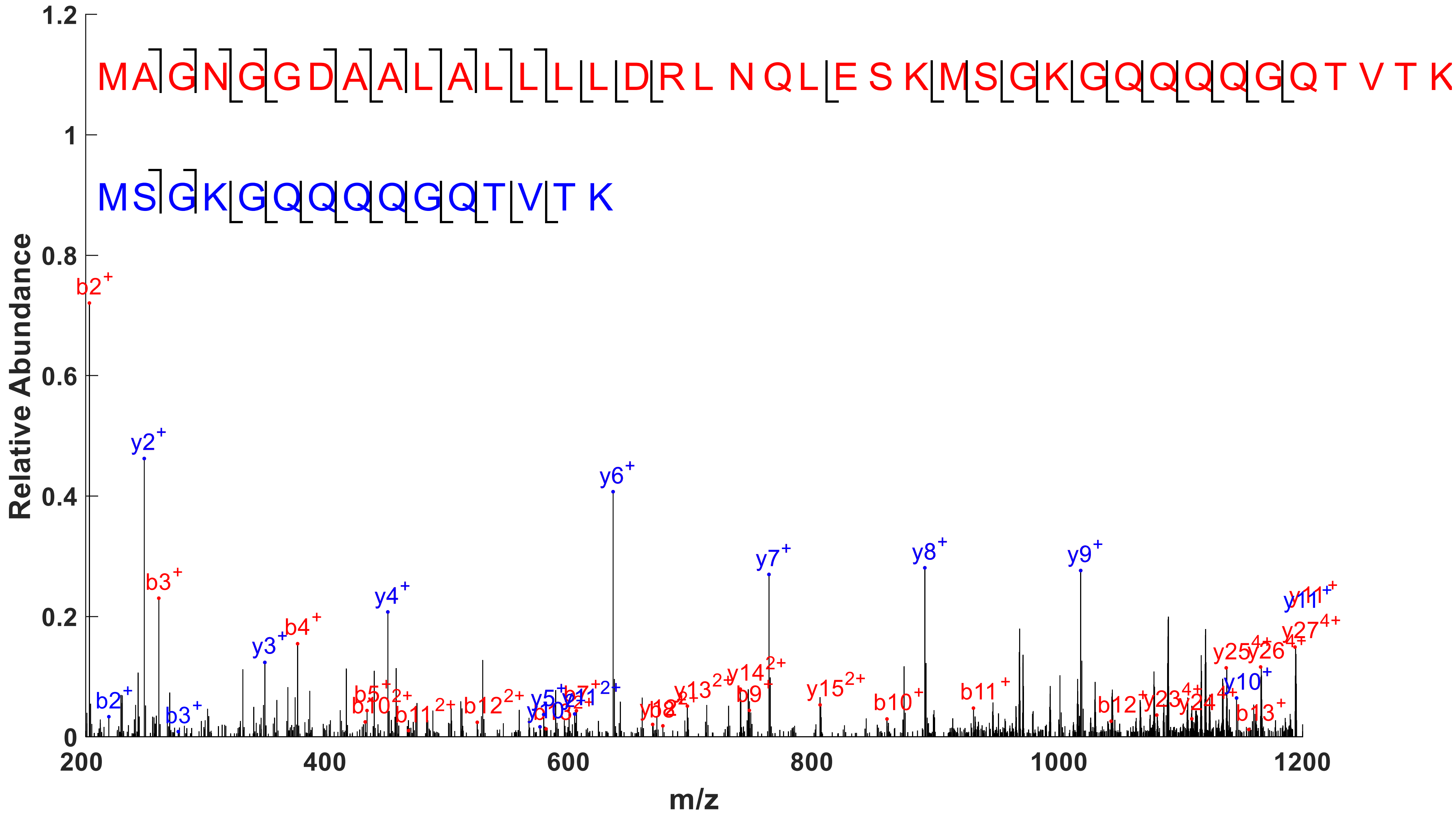


Protein N dimer cross link (4) K233 – K237.

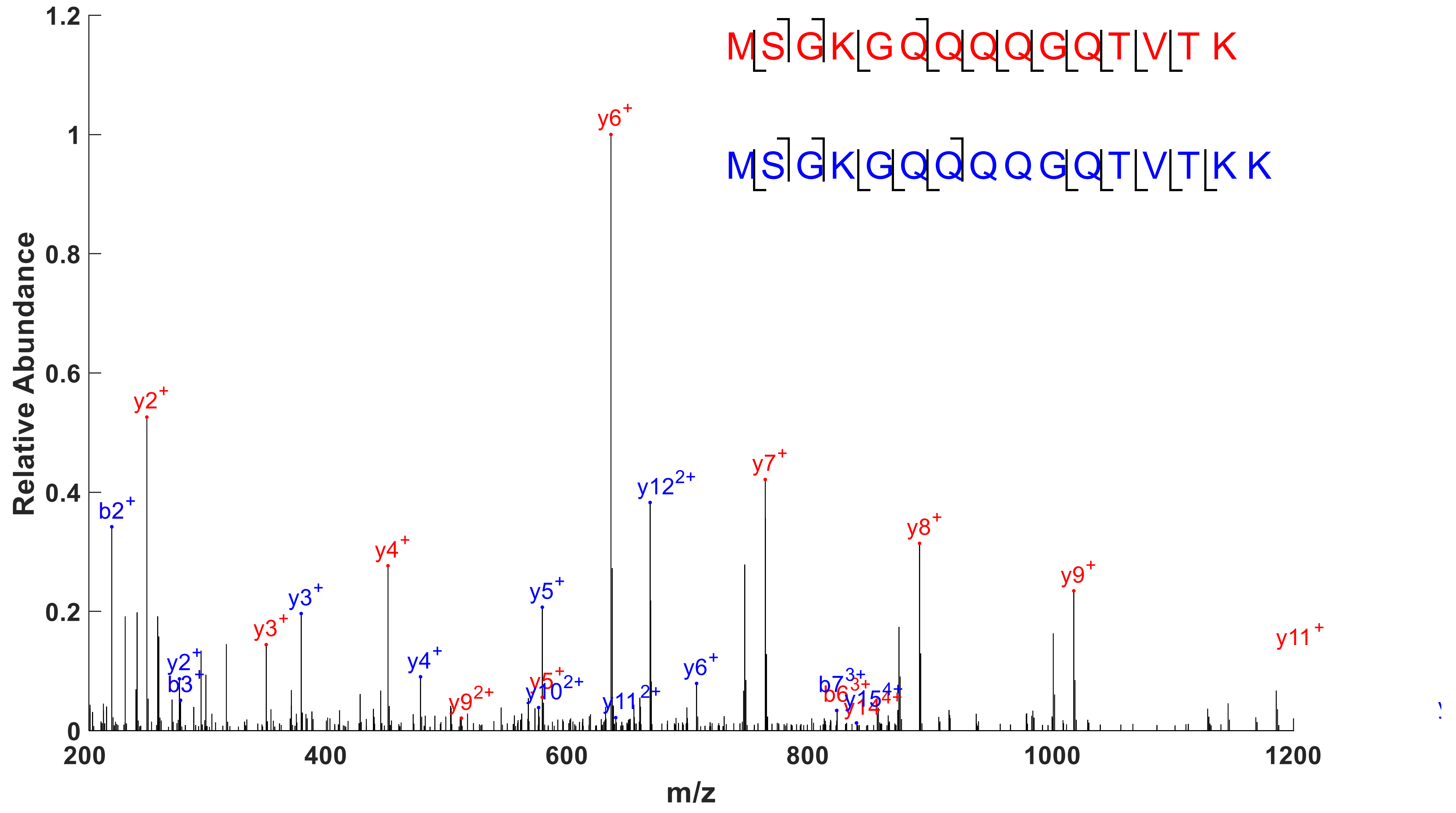


Protein N dimer cross link (5) K237 – K237.

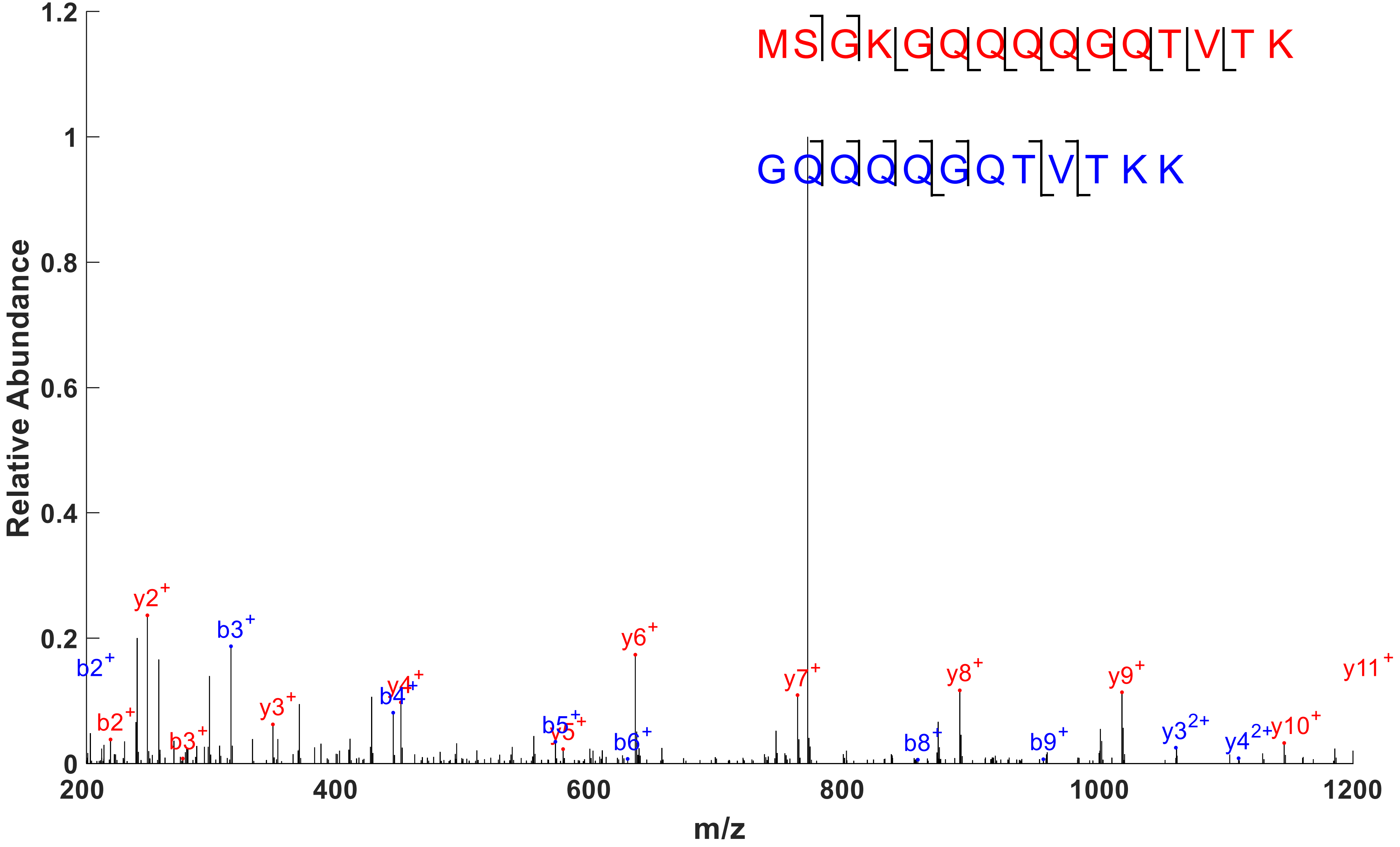


Protein N dimer cross link (6) K237 – K248.

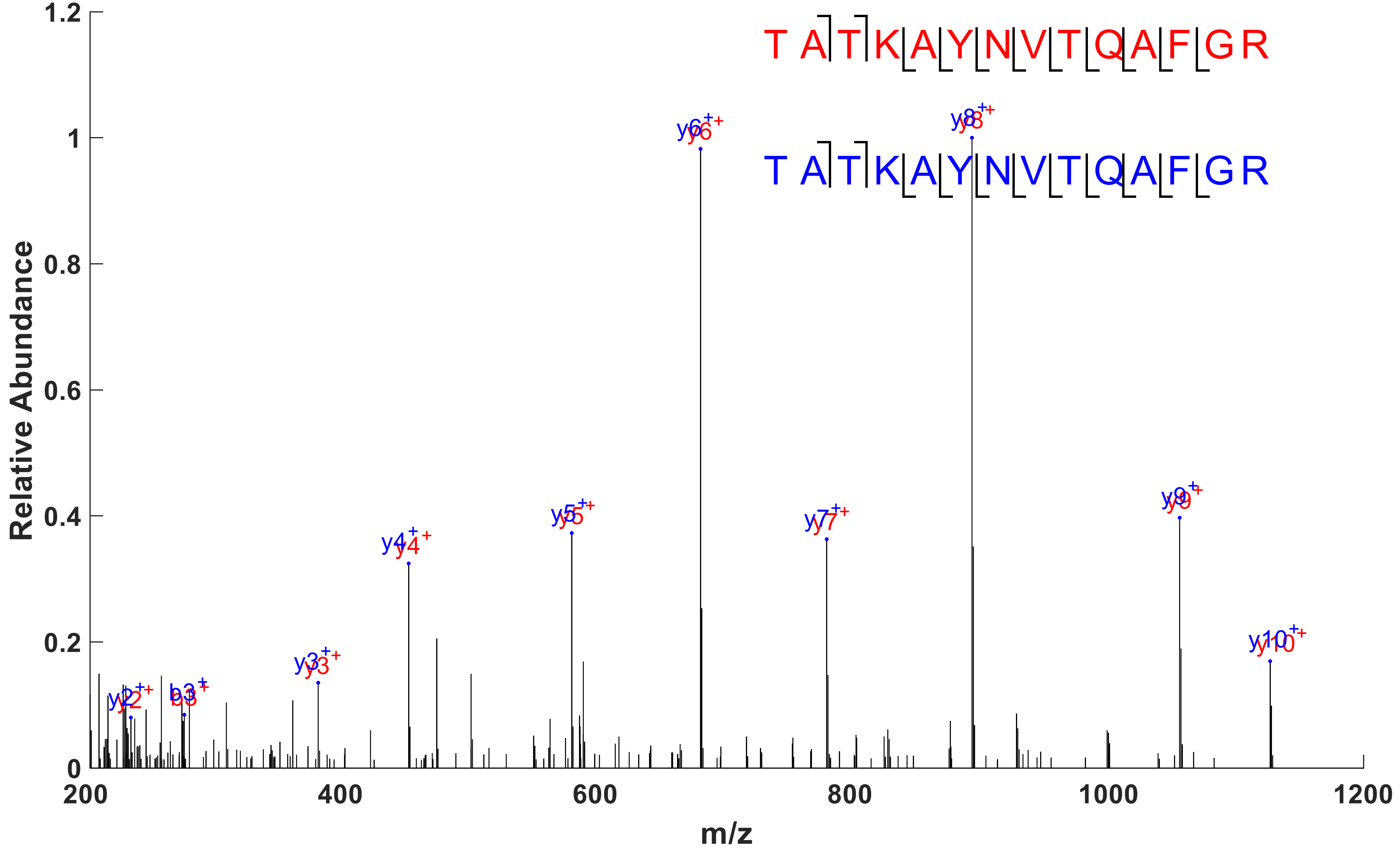


Protein N dimer cross link (7) K266 – K266.

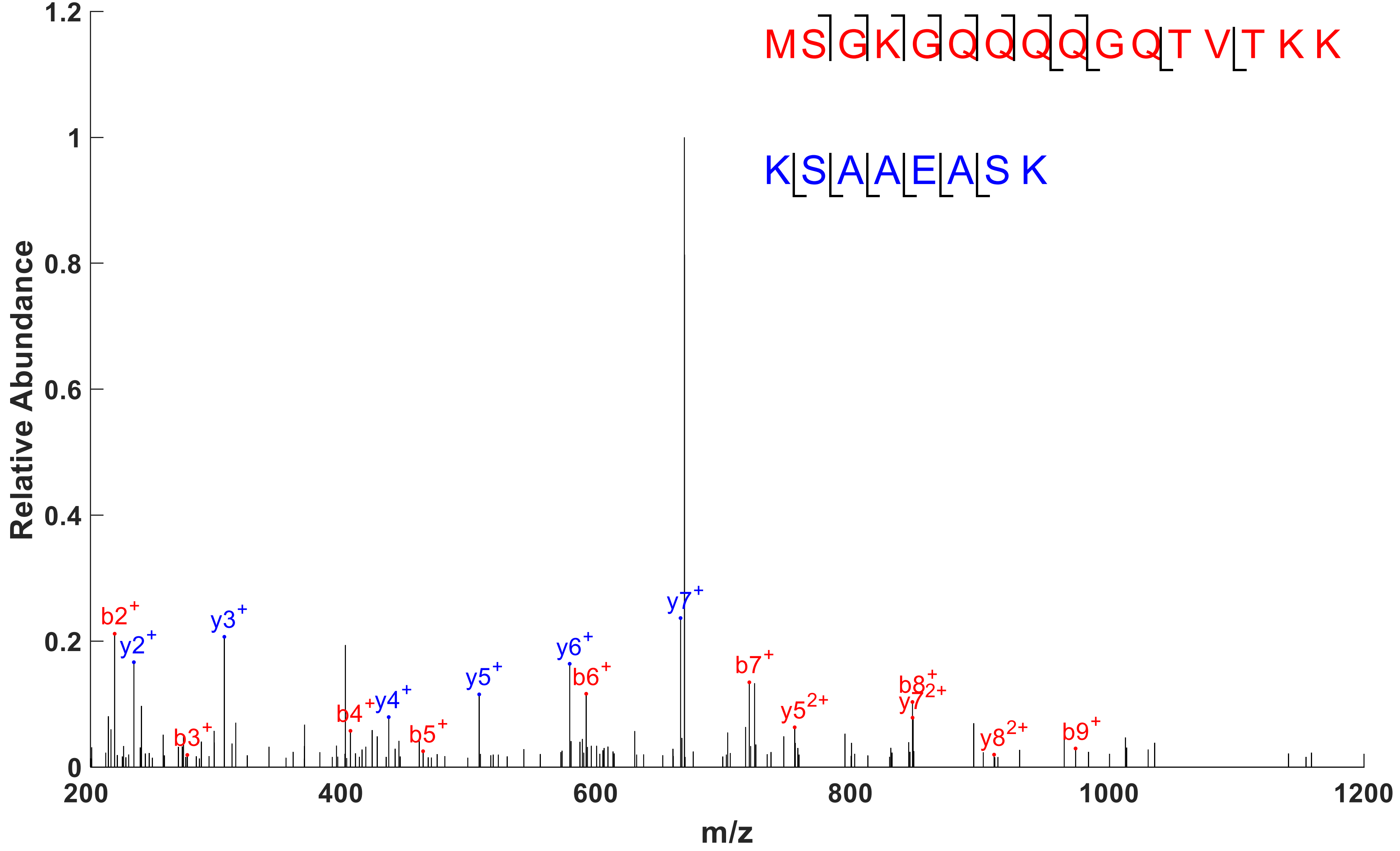


Protein N dimer cross link (8) K248 – K249.

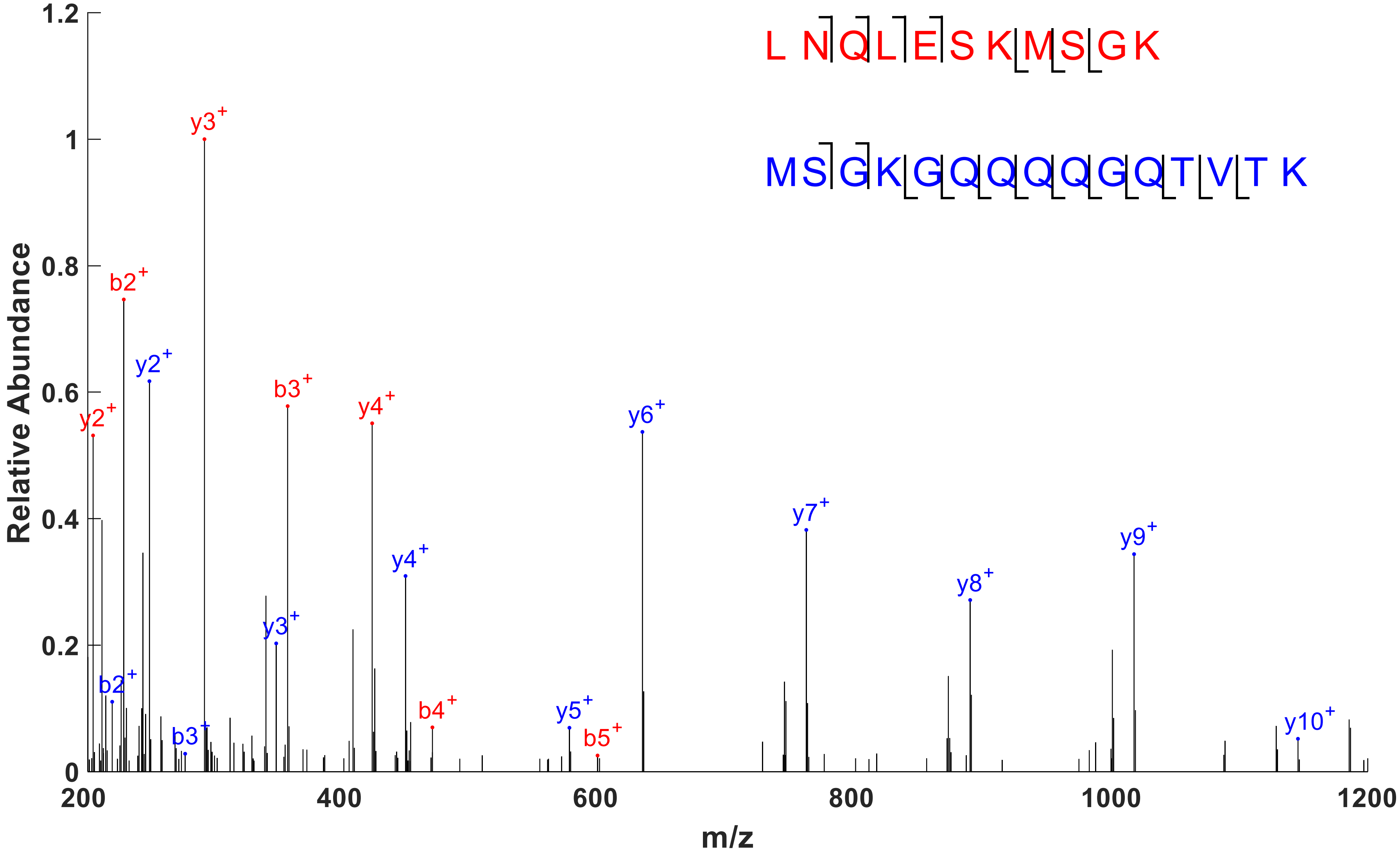


Protein N dimer cross link (9) K233 – K237.

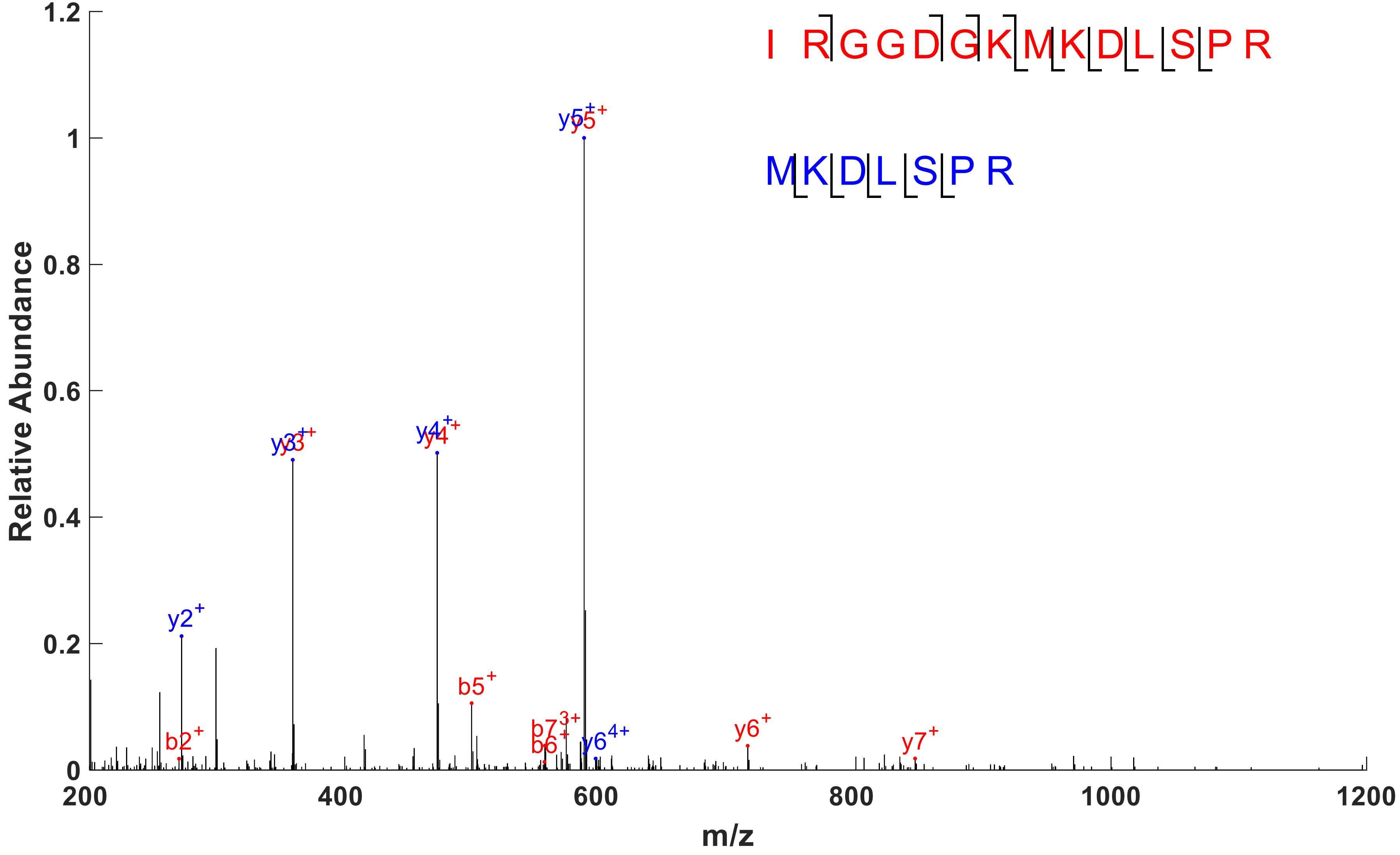


Protein N dimer cross link (10) K100 – K102.
